# Tissue-guided multi-omics profiling identifies extracellular vesicle biomarkers indicative of lung pathology in acute respiratory distress syndrome

**DOI:** 10.64898/2025.12.09.693331

**Authors:** Po-Tsang Chen, Chao-Yuan Chang, Chi-Li Chung, Chih-Hsin Lee, Chi-Won Suk, Chiou-Feng Lin, Yu-Jui Fan, Yen-Wen Lu, Yi-Chiung Hsu, Tanyu Chang, Chun-Jen Huang, I-Lin Tsai

**Author notes:** Corresponding authors: **Yi-Chiung Hsu*, Ph.D.**, Department of Biomedical Sciences and Engineering, National Central University, Taoyuan City, Taiwan,; **Chun-Jen Huang*, M.D., Ph.D.**, Graduate Institute of Clinical Medicine, College of Medicine, Taipei Medical University, Taipei, Taiwan,; **I-Lin Tsai*, Ph.D.**, Department of Biochemistry and Molecular Cell Biology, School of Medicine, College of Medicine, Taipei Medical University, Taipei, Taiwan.

## Abstract

**Background:** Acute respiratory distress syndrome (ARDS) remains a lethal inflammatory lung condition lacking reliable biomarkers that reflect lung-specific pathology. Extracellular vesicles (EVs) circulate systemically and may carry molecular signals from injured organs, but the correspondence between EV cargo and lung tissue alterations remains unclear.

**Methods:** We established aspiration-, lipopolysaccharide (LPS)-, and COVID-19–induced murine ARDS models and applied a tissue-guided multiomics framework integrating proteomic and metabolomic analyses of lung tissue and plasma-derived EVs to identify lung-originating circulating biomarkers.

**Results:** Four proteins—haptoglobin (HP), inter-alpha-trypsin inhibitor heavy chains 3 and 4 (ITIH3, ITIH4), and clusterin (CLU)—were consistently upregulated in both lung tissue and plasma EVs across all ARDS etiologies. Metabolomic integration revealed dysregulation of arachidonic acid metabolism as a unifying inflammatory axis. Multiomics network analysis further distinguished etiology-specific molecular programs, including glycolytic activation in aspiration-induced, platelet aggregation in LPS-induced, and vascular smooth muscle dysregulation in COVID-19–induced ARDS.

**Conclusions:** This study establishes a tissue-informed EV profiling framework that links local lung pathology to systemic molecular signatures, revealing HP, ITIH3, ITIH4, CLU, and arachidonic-acid–related metabolites as potential diagnostic markers for ARDS. These findings provide a foundation for developing clinically translatable, EV-based biomarker assays for early detection and molecular subtyping of lung injury.

## 1. BACKGROUND

Acute respiratory distress syndrome (ARDS) remains a devastating clinical condition marked by diffuse alveolar injury, systemic inflammation, and high mortality despite decades of research [1]. It can arise from diverse pulmonary or extrapulmonary insults such as aspiration, sepsis, or viral infection, leading to widespread inflammation and disruption of the alveolar–capillary barrier [2–4]. Current diagnostic criteria rely largely on physiological parameters and imaging, which fail to capture the underlying molecular pathology. As a result, there is still no reliable biomarker that reflects the dynamic state of lung injury or guides precision management. [5–7].

Extracellular vesicles (EVs) have recently gained attention as promising candidates for biomarker discovery in inflammatory and pulmonary diseases [8–10]. EVs are nano-sized, membrane-bound vesicles released by most cell types and carry proteins, lipids, and nucleic acids that mirror the physiological or pathological states of the origin cells. Because plasma-derived EVs circulate systemically, translational application of EVs for diagnostic and prognostic applications can be important. However, plasma EVs originate from multiple tissues and cell types, making it difficult to discern which molecular signals truly reflect pulmonary pathology [11, 12].

To overcome this limitation, integrating molecular information from lung tissue can and plasma EV data can help to define specific molecular signatures that originate from lungs. This tissue-guided strategy links local injury with systemic responses and provides a biologically grounded framework for EV-based biomarker discovery.

Despite growing interest in this field, few studies have systematically compared the proteomic and metabolomic profiles of EVs of various etiologies of lung injury. In this study, we established aspiration-induced, lipopolysaccharide (LPS)-induced, and COVID-19-induced ARDS murine models to capture distinct etiologies of lung injury. Using integrated multiomics analyses of lung tissue and plasma EVs, we aimed to identify both shared molecular features and tissue-guided plasma biomarkers that could be indicative of the lung pathology. This approach provided new insights into the molecular landscape of ARDS and advanced the development of biologically informed, non-invasive biomarker discovery [13–17].

## 2. METHODS

### 2.1 Collections and Preparation of ARDS Mouse Models

#### 2.1.1 Animals and animal care

All mouse models were provided by the laboratory of Professor Chun-Jen Huang at the Graduate Institute of Clinical Medicine, Taipei Medical University. The Institutional Animal Use and Care Committee of Taipei Medical University approved all animal studies (LAC-2022-0427). Adult male wild-type C57B/L6 mice (7–8 weeks old, Taiwan National Laboratory Animal Center, Taipei, Taiwan) were used for the experiments. Regular laboratory chow and water were provided to all mice with free access. Mice were maintained on a 12:12 h light–dark cycle routine. Care and handling of the mice were performed in accordance with the US National Institutes of Health guidelines.

#### 2.1.2 ARDS mouse models

Sepsis-induced ARDS model was generated using intraperitoneal injection of gram (-) endotoxin (25 mg/kg, lipopolysaccharide, LPS, Escherichia coli 0127:B8, Sigma-Aldrich, USA), as previously reported by us [18]. Aspiration pneumonia (AP)-induced ARDS model was generated using previous described protocols [19]. A gastric-content mimic (50 μL) containing xanthan gum-based thickener (12 mg/mL, ThickenUp®; Nestlé Health Science), pepsin (2 mg/mL; Sigma–Aldrich), and lipopolysaccharide (2.5 mg/mL; Escherichia coli 0127:B8, Sigma–Aldrich), adjusted to pH 1.6 with HCl (Sigma–Aldrich), was administered via oropharyngeal instillation to anesthetized mice (anesthetized with 3% isoflurane) to simulate macroaspiration.

The COVID-19 mimic-induced ARDS model was generated through intratracheal administration of the SARS-CoV-2 spike protein S1 subunit (15 μg/mouse; RayBiotech, USA) plus LPS (1 mg/mouse; Sigma-Aldrich), according to previous protocols [20] (Supplementary Fig. S1).

This experiment used an independent cohort of 55 mice. From each group, 6 mice underwent lung injury assessment and tissue sectioning. Among these, 6 mice were assigned to the sham group, 7 of them were assigned to the LPS group, and 18 of them were assigned to the AP and COVID-19 groups (n = 9 in each group).

At 48 h after ARDS induction, all surviving mice were anesthetized (Zoletil/xylazine, 40/10 mg/kg, i.p.) and blood samples obtained via cardiac puncture. These blood samples were then placed in heparin tubes (Venosafe; Terumo Europe, Leuven, Belgium) and centrifuged at 2000 × g for 10 min. The resulting supernatant plasma samples were collected and stored at −20 °C for subsequent analysis.

Following euthanasia by decapitation, the lung tissues of the mice from each group were removed en bloc and promptly snap-frozen in liquid nitrogen, and preserved at −80 °C for subsequent analysis.

#### 2.1.3 Lung injury evaluation

An independent cohort of 55 mice was used for this lung injury evaluation. In this cohort, 12 mice were allocated to a sham group, 13 mice LPS group, 30 mice were allocated to the AP and COVID-19 groups (n = 15 in each group). Sample size of each group was determined based on the 48-h survival rate data, to ensure each group would have at least 6 surviving mice for this assay. At 48 h after LPS or saline injection, 6 surviving mice from each group were anesthetized (zoletil/xylazine, 40/10 mg/kg, i.p.). Subsequently, tracheostomy was done on the mice to insert tracheostomy tube (22# intravenous catheter; Terumo Corp., Tokyo, Japan) to facilitate pulmonary function assay using a computerized small animal ventilator (flexiVent FX; SCIREQ Inc., Montreal, QC, Canada). The mechanical ventilation was set at a ventilation rate of 150 breaths/min and a tidal volume of 0.2 mL. Parameters including inspiratory capacity, airway resistance, and dynamic compliance, were recorded using a FlexiWare 8 System (SCIREQ) [21].

Formaldehyde-infused right lung tissues from 6 surviving mice in each group were embedded in paraffin wax, serial sectioned, and then stained with hematoxylin and eosin [22].

#### 2.1.4 Lung tissue sample preparation

At 48 h following euthanasia by decapitation, lung tissues of the first subset of 6 surviving mice from each group were removed en bloc and promptly snap-frozen in liquid nitrogen, preserving them at −80 °C for subsequent analysis.

Lung tissues from four ARDS mouse groups were cut on ice to <1×1×5□mm, followed by 50□mg of tissue being mixed with equal-weight beads and 500□µL RIPA buffer. Samples were homogenized using a bullet blender with stepwise speed settings, cooling between cycles, and then centrifuged (5g, 5□min) to collect the supernatant.

### 2.2 Extracellular vesicles

#### 2.2.1 Isolation and purification

Before isolating EVs from mouse plasma, samples were thawed on ice and centrifuged at 3000g for 15 minutes to remove cell debris, followed by centrifugation at 12000g for 15 minutes to eliminate large vesicles. The supernatant was then processed using a SmartSEC single column. After removing the storage buffer and washing the resin with isolation buffer, 250□μL of plasma was loaded into the column, followed by 250□μL of isolation buffer. The column was incubated for 30 minutes at room temperature with mixing using a rotator and vortex mixer. EVs were then eluted by centrifugation at 500g for 30 seconds and pooled for storage at 4°C.

#### 2.2.2 Characterization of extracellular vesicles

To confirm the presence and characteristics of EVs, we used western blotting, nanoparticle tracking analysis (NTA), and transmission electron microscopy (TEM). Western blotting was used to detect exosomal markers CD63, CD81, and CD9. EV proteins were extracted using RIPA buffer, sonicated, denatured, and separated by SDS-PAGE. After transferring to a PVDF membrane, specific bands were probed with primary and HRP-conjugated secondary antibodies, and visualized using chemiluminescence. NTA was conducted using NanoSight NS300 to assess particle size and concentration. EVs were diluted in PBS, and a 2500-fold dilution provided optimal measurement clarity. TEM analysis was performed to visualize EV morphology. Samples were stained with 2% uranyl acetate and observed under a Hitachi HT-7700 microscope. Together, these methods confirmed the identity, size distribution, and structural integrity of the isolated EVs.

### 2.3 Proteomics analysis

#### 2.3.1 Proteomics sample preparation and parameters for LC–MS/MS analysis

To ensure equal protein amounts for mass spectrometry (MS), we quantified protein concentrations using a Pierce™ BCA protein assay kit. BSA standards (25–2000 μg/mL) were prepared for the calibration curve and the samples were mixed with RIPA buffer, sonicated, incubated with BCA working reagent (solvent A/B = 50:1) Reading were measured at 37°C using an ELISA reader. Samples that exceeded the curve range were diluted appropriately. For protein cleanup, 50 μg of protein was precipitated using cold acetone (4:1 ratio), incubated at –20°C overnight, and centrifuged. The pellet was washed, redissolved in 6M urea, reduced with 550 mM DTT, alkylated with 450 mM IAA, and digested overnight with trypsin. Digestion was stopped with formic acid. For final purification, solid-phase extraction (SPE) was performed using methanol, formic acid, and acetonitrile solvents to remove salts and concentrate analytes. Eluted samples were dried with SpeedVac and stored at –20°C until MS analysis.

Details on proteomics material, LC–MS/MS analysis, and parameter settings are provided in the Supporting Information.

#### 2.3.2 Data Analysis and identification for proteomics

Raw MS data from proteomics were analyzed using MaxQuant with LFQ enabled, trypsin digestion, and specific modifications. For global parameters, we have uploaded a mice fasta file (uniprotkb_taxonomy_id_10090_AND_reviewed_2023_10_23.fasta) from UniProt since we were analyzing the human proteome. As for the identification, we switched ‘match between runs’ to scan for the missing MS1 features across the retention time plane between runs.

#### 2.3.3 Proteomics data cleaning

Data were cleant in Perseus by removing contaminants and filtering for valid intensity values. We filtered out ‘reverse’, ‘only identified by site’, and ‘potential contaminant’ proteins. Then we created four groups, i.e., control, AP, LPS, and COVID-19 groups. Next, we filtered rows to obtain at least 60% of valid total intensity values, which meant the 60 % of the values inside the rows should not be NaN. Finally, we renamed the column names and exported the data as a text (.txt) file.

#### 2.3.4. Screening of differentially expressed proteins and functional enrichment data

MetaboAnalyst 6.0 was used to conduct downstream analyses, including hierarchical clustering, sPLS-DA, and volcano plots. Differentially expressed proteins (DEPs) are defined as proteins with a Wilcoxon test p-value < 0.05 and fold change > 1.3.Gene enrichment study of the exosomal proteome was performed using FunRich software (v3.1.3). To conduct gene enrichment analysis, we input the total proteins of lung tissue and plasma-EV dataset (the proteins identified in each dataset). Subsequently, the UniProt accession number of total proteins was converted to genes that were mapped in the database. A functional analysis was performed. Proteins were classified by cellular component, molecular function, and biological process. The enriched GO terms were screened based on p value >0.05 and arranged according to the number of proteins.

### 2.4 Metabolomics analysis

#### 2.4.1 Metabolomics sample preparation

From acetone obtained in the previous step, a vacuum centrifugal concentrator (SpeedVac) was used to dry the samples. After drying, 50 µL of methanol was added to re-dissolve the precipitate. Then, the solution was sonicated in a water bath for 15 minutes.

Next, 50 µL of water was added and sonicated for another 15 minutes. Finally, the mixture was centrifuged at 12,000 g for 10 minutes. After centrifugation, 60 µL of the supernatant from each sample was removed, ensuring that the precipitate was not disturbed to avoid interference with the mass spectrometer operations. Details of metabolomics meterial, LC–MS/MS analysis, and parameter settings are provided in the Supporting Information.

#### 2.4.2 Metabolomics data analysis and identification

Metabolomics data from Progenesis QI measurement files were filtered to retain rows with at least 60% valid intensity values.

Mass spectrometry analysis was performed on lung tissue metabolomics samples, excluding COVID-19 3 and COVID-19 4 due to instrument issues, resulting in seven COVID samples for analysis. Raw data from metabolomics were uploaded to Progenesis QI using centroided data at 20,000 FWHM resolution. Adducts were defined for both positive and negative ion modes and data were aligned using Waters (.raw) files. After grouping samples intos, LPS, AP, and COVID-19 groups, peak picking was conducted and compound identification was done using MetaScope with HMDB SDF files (5 ppm tolerance). We chose Waters’ search method, MetaScope in the identify compounds section for its flexibility in searching compound data. We uploaded a Metabolite Structures SDF file from the Human Metabolome Database (HMDB) to serve as our compound database. We set both the precursor and fragment tolerances to 5 ppm for identifying compounds. Unidentified features were removed, and we selected possible adducts for our positively and negatively charged modifications including metabolites with specific adducts. ([M+H]^+^, [M+Na]+, [M+K]^+^, [M+2H]²□, [M–H]□)

#### 2.4.3. Screening of differentially expressed metabolites and pathway analysis

The ARDS mouse model lung tissue and plasma-EV values dataset was obtained. Next, we used MetaboAnalyst 6.0 to analyze the differential expression features (DEFs) of each of the three ARDS groups compared with the sham group. For the volcano plot and heatmap analyses, sample normalization was performed using median values. Raw *p* value threshold was set to 0.05. Data was analyzed using sPLS-DA to interpret group differences with the sample being normalized based on median values and data auto-scaling. The acceptable mass error range was set between -5 and 5 ppm. Furthermore, only features with an isotope similarity greater than 90% and a Progenesis score exceeding 36 were retained. After filtering based on these parameters, the corresponding HMDB IDs were extracted and used for downstream analyses.

A pathway analysis was done using MetaboAnalyst 6.0. The first step of differential expression screening was done on the positively charged and negatively charged compound identification data in the metabolomics data. Finally, the HMDB IDs identified by MetaboAnalyst ID were compared with MetaboAnalyst built-in KEGG database pathway analysis to identify the final number of features.

### 2.5 Statistical analysis

To identify potential ARDS biomarkers, we analyzed LFQ data from lung tissue and plasma-EV using MetaboAnalyst to screen for DEPs (FC > 1.3, *p* < 0.05). This was followed by Mann-Whitney tests (using Prism 8) for validation. Proteins and features that were consistently upregulated or downregulated in both sample types were further examined through box plots for potential biomarkers.

### 2.6 Multiomics analysis

Lung tissue samples from three types of ARDS mouse models underwent integrated proteomic and metabolomic analysis using OmicsNet 2.0. Lung tissue data included DEPs and DEMs with log2(FC) values, and all networks were constructed using KEGG-derived protein-metabolite interactions. Subnetworks were generated based on these interactions, and over-representation analysis was performed on all nodes (proteins and metabolites) against the KEGG database.

## 3. RESULTS

### 3.1 Establishment of ARDS mouse models and extracellular vesicle characterization

The overall study design is illustrated in Figure 1. Lung injury was evaluated using formaldehyde-perfused right lung tissue from 6 mice per group. Histological examination (Figure 2A) revealed normal alveolar structures in sham mice, whereas the LPS-, AP-, and COVID-19-induced groups exhibited typical ARDS features, including alveolar wall thickening and inflammatory infiltration. Corresponding lung injury scores were significantly higher in all ARDS groups compared with sham controls (*p* < 0.01). Pulmonary function tests (Figure 2B,C) further demonstrated marked impairment with reduced inspiratory volume, dynamic compliance, and vital capacity, accompanied by increased airway resistance and elastance (*p* < 0.01).

**Figure 1.**
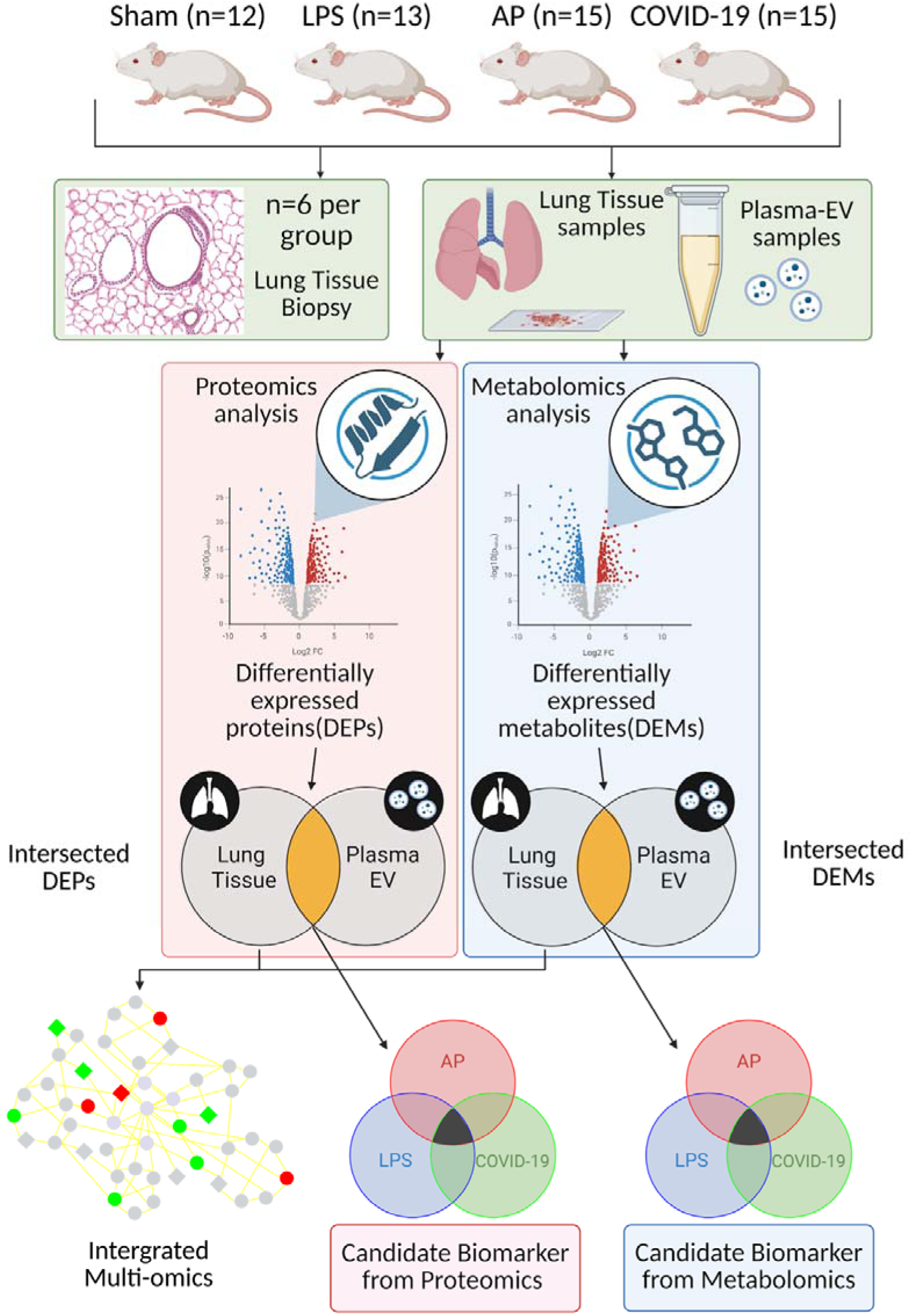
Flow chart of study design.

**Figure 2.**
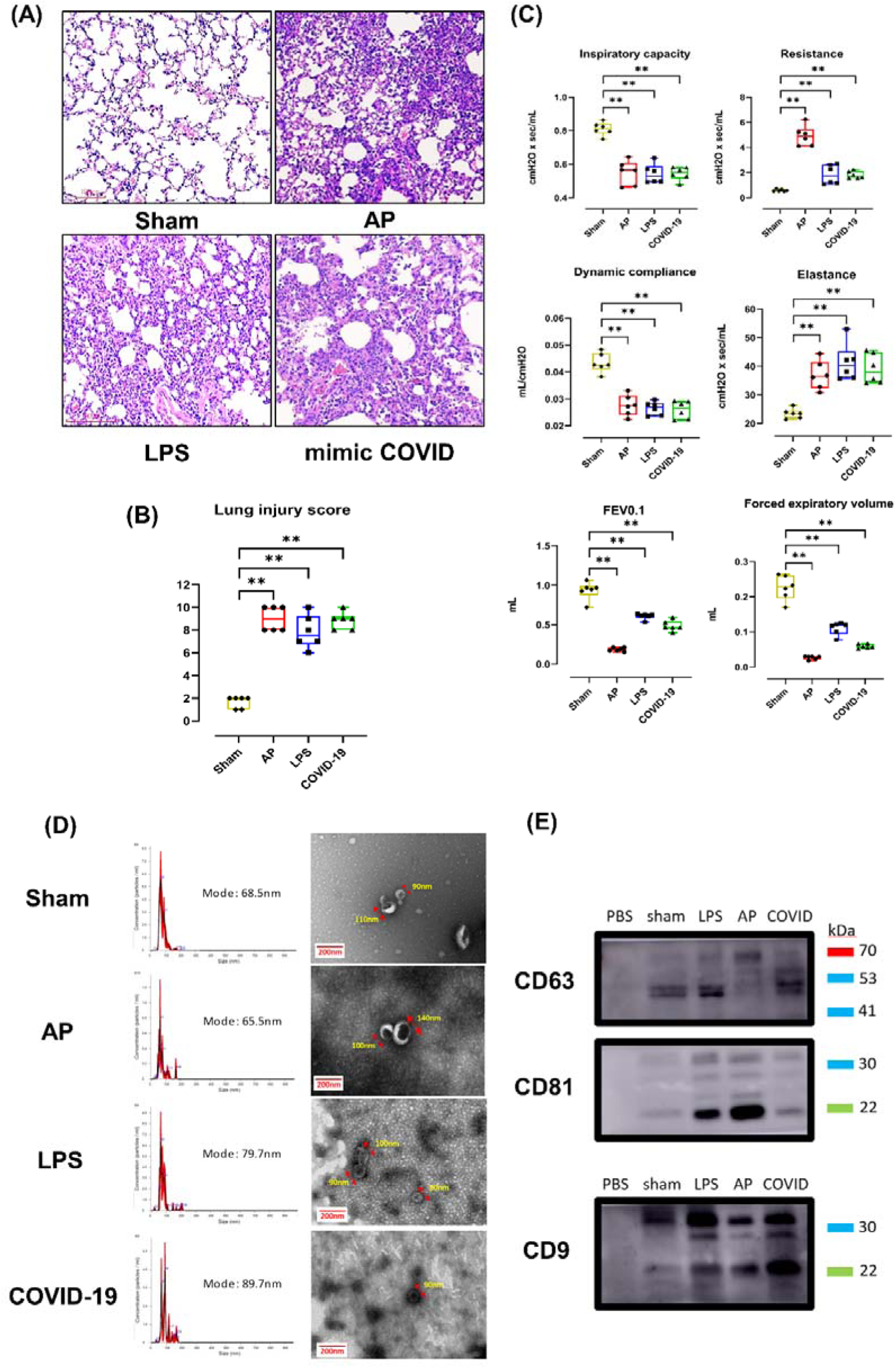
Establishment of ARDS mouse models and extracellular vesicle characterization. Histological examination of four groups of the lung tissue from ARDS mouse model. (B-C) Box plot analysis the lung injury score, inspiratory capacity, airway resistance, and dynamic compliance comparing in the three ARDS models with the Sham group. The Mann-Whitney test was conducted after ruling out the outliers. The p values were presented with asterisks (ns: P > 0.05, *: P ≤ 0.05, **: P ≤ 0.01, ***: P ≤ 0.001, ****: P ≤ 0.0001). (D) Representative nanoparticle tracking analysis and morphology of plasma-EVs from four groups of ARDS model mice under TEM. Scale bars, 200 nm. (E) The expressions of CD63, CD81, and CD9 in the plasma-EV were analyzed using Western blot.

Isolated EVs displayed the expected cup-shaped morphology and bilayer membrane (Figure 2D), with particle diameters ranging from 30 to 150 nm. Western blotting confirmed the presence of EV surface markers CD9, CD63, and CD81 (Figure 2E). Together, these findings validated the successful establishment of ARDS models and the consistent isolation of plasma-derived EVs suitable for subsequent omics analyses.

### 3.2 Proteomic profiling of lung tissue and plasma extracellular vesicles

sPLS-DA analysis revealed clear separation between sham and ARDS groups in both datasets, with component 1 explaining 22.8% and 19.7% of the variance, respectively (Figure 3A, D). Heatmap visualization confirmed high within-group reproducibility (Figure 3B, E). In the lung tissue dataset, volcano plot analyses identified 577, 281, and 204 DEPs in the AP, LPS, and COVID-19 groups comparing with sham control, respectively (Figure S2). Among these, 125 were shared across all ARDS models (98 upregulated and 26 downregulated) (Figure 3C, Table S1). In the plasma EV dataset, 88, 80, 38 DEPs were identified in the AP, LPS, and COVID-19 groups compared with the sham control, respectively (Figure S3). Among these, 10 were shared across all ARDS models (7 upregulated and 3 downregulated) (Figure 3F).,

**Figure 3.**
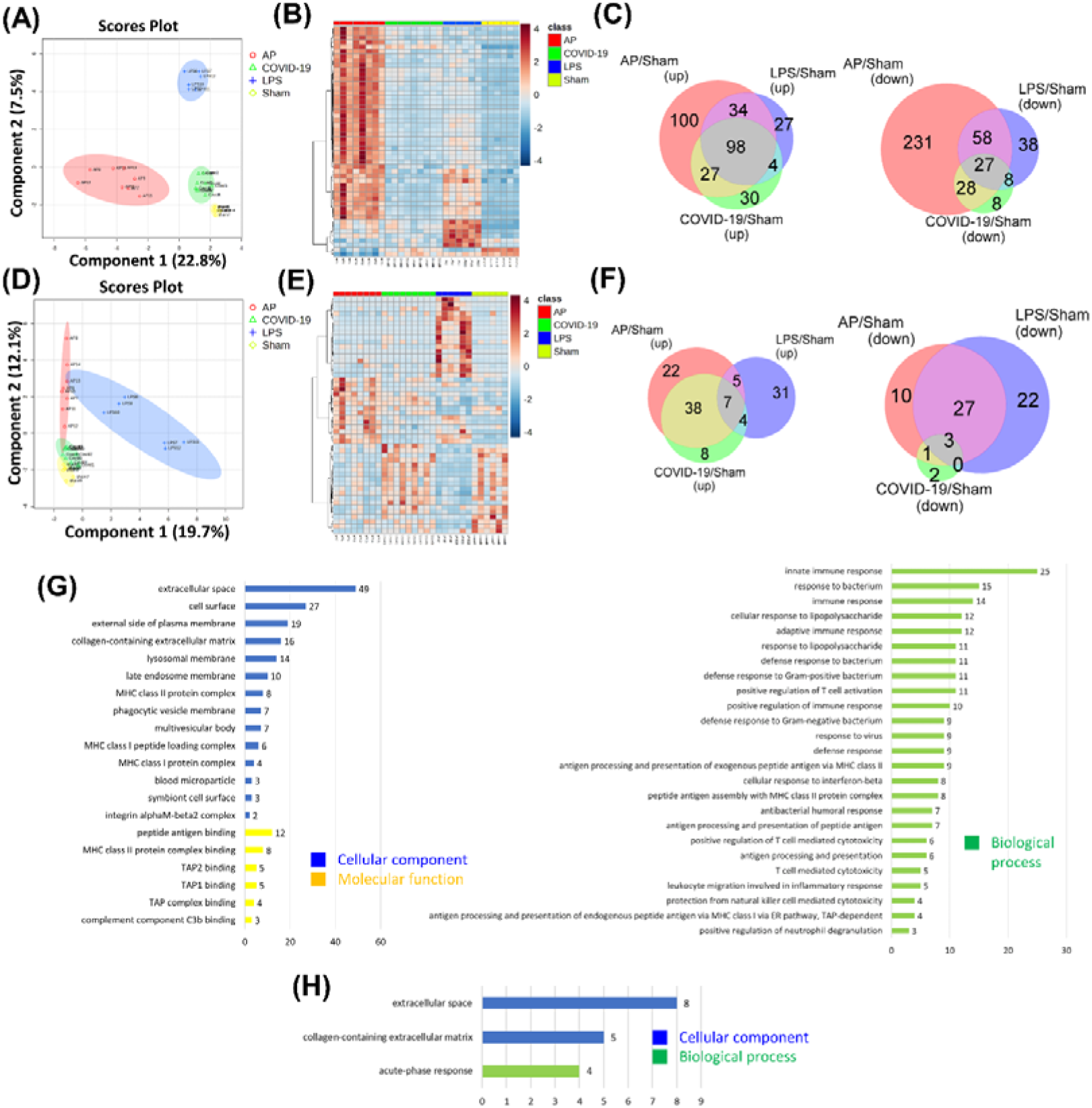
Proteomics data from lung tissue and plasmaEV samples with ARDS mouse groups. (A)Score plot of sPLS-DA analysis showing the distribution between sham group, LPS group, AP group, and COVID-19 group’s lung tissue sample within a 95% confidence region. (B) heatmap plot showing the top 50 protein abundance between the sham group, LPS group, AP group, and COVID-19 group with lung tissue samples. (C) Venn diagrams illustrating the overlap among differentially expressed the lung tissue proteins from the three ARDS models, significantly increased proteins (left), and significantly decreased proteins (right). (D)Score plot of sPLS-DA analysis showing the distribution between sham group, LPS group, AP group, and COVID-19 group’s plasma-EV sample within a 95% confidence region. (E) heatmap plot showing the top 50 protein abundance between the sham group, LPS group, AP group, and COVID-19 group with plasma-EV samples. (F) Venn diagrams illustrating the overlap among differentially expressed the plasma-EV proteins from the three ARDS models, significantly increased proteins (left), and significantly decreased proteins (right). The bar plots on the right depict enriched Gene Ontology (GO) terms in cellular component, molecular function, and biological process categories for the overlapping DEPs. (G) Gene Ontology-based functional classification of differentially abundant lung tissue proteins. (H) Gene Ontology-based functional classification of differentially abundant plasma-EV proteins.

Gene ontology enrichment analysis indicated that the common DEPs in the lung tissue dataset were primarily associated with extracellular space, peptide antigen binding, and innate immune response pathways (Figure 3G), whereas in the plasma EV dataset, the common DEPs enriched in extracellular matrix components and acute phase response pathways (Figure 3H). Notably, serum amyloid A1 was consistently upregulated in both lung tissue and plasma-EV datasets in AP and COVID-19 models compared with sham controls (Figure S5).

### 3.3 Comparative metabolomic signatures across lung tissue and plasma extracellular vesicles

The overall workflow for metabolomic data acquisition and feature identification is summarized in Figure S6. sPLS-DA analysis revealed distinct separation between sham and ARDS groups in both positive and negative ionization modes, with well-defined clustering among the LPS, AP, and COVID-19 models for both datasets (Figure 4A–H). Volcano plots identified 1951 (1272 positive; 679 negative), 1237 (1070 positive; 167 negative), and 1127 (851 positive; 276 negative) DEFs in the AP, LPS, and COVID-19 groups for the lung tissue dataset, respectively (Supplementary Figs. S7–S8).

**Figure 4.**
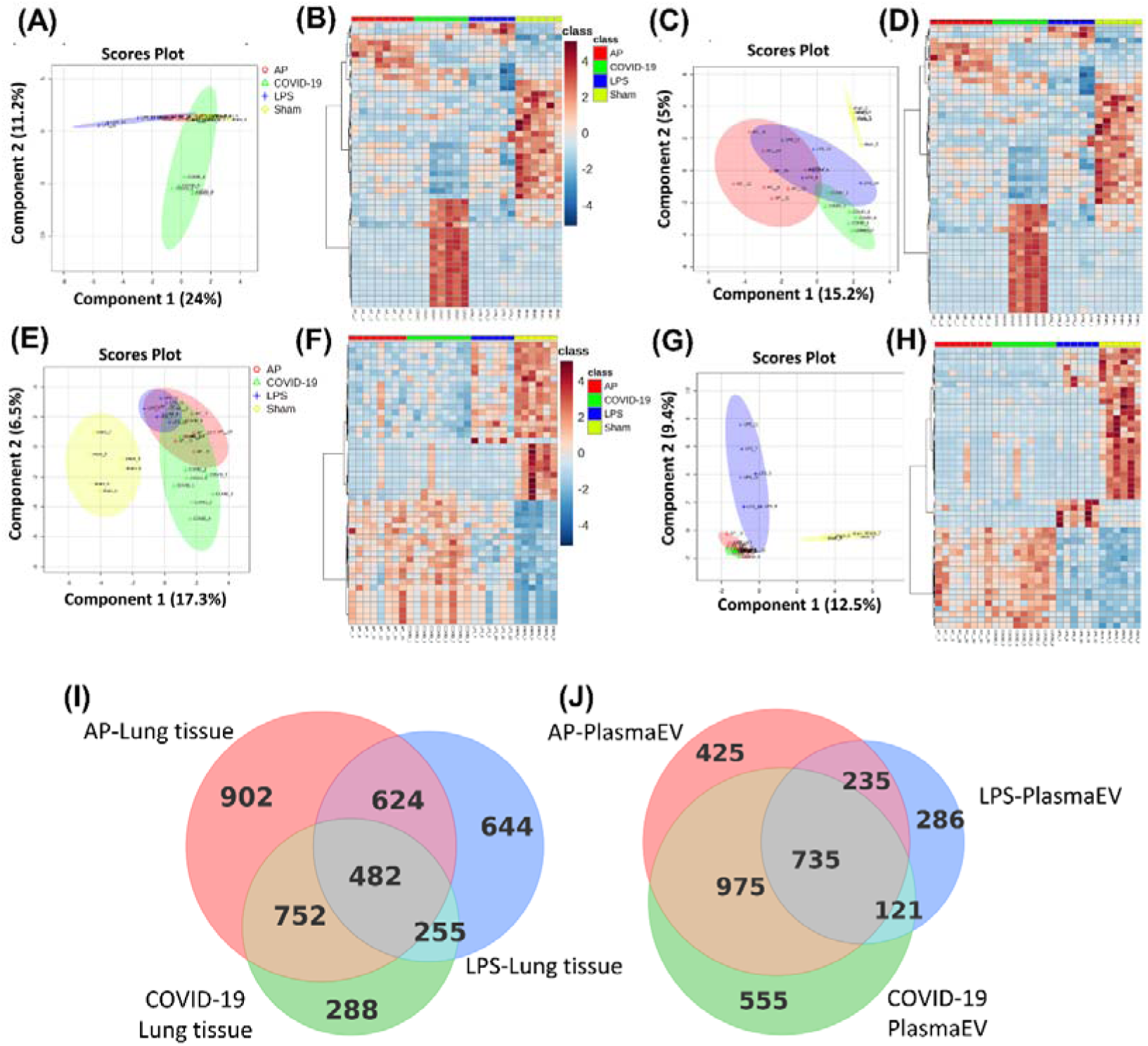
Metabolomics data from lung tissue and plasmaEV samples with ARDS mouse groups. (A, C) Score plots of sPLS-DA showing the separation among the sham, LPS, AP, and COVID-19 groups of lung tissue samples in both positive and negative ion modes within the 95% confidence region.(B, D) heatmap plots showing the top 50 protein abundance between the sham group, LPS group, AP group, and COVID-19 group with lung tissue samples in the positive and negative mode.(E, G) Score plots of sPLS-DA showing the separation among the sham, LPS, AP, and COVID-19 groups of plasma-EV samples in both positive and negative ion modes within the 95% confidence region.(F, H) heatmap plots showing the top 50 protein abundance between the sham group, LPS group, AP group, and COVID-19 group with plasma-EV samples in the positive and negative mode.(I) Venn diagrams illustrating the overlap among differentially expressed lung tissue proteins from the three ARDS models. (J) Venn diagrams illustrating the overlap among differentially expressed plasma-EV proteins from the three ARDS models.

As for the plasma EVs dataset, a total of 2308 (1047 positive; 1261 negative), 1074 (498 positive; 576 negative), and 2606 (1346 positive; 1260 negative) DEFs were identified, respectively (Supplementary Figs. S9–S10). Comparative analysis revealed 482 shared DEMs in the lung tissue datasets and 735 shared DEMs in plasma-EVs (Figure 4I,4J).

### 3.4 Tissue-guided correlation between lung and plasma EV proteomes

To explore the molecular correspondence between lung tissue and circulating EVs, we intersected DEPs from both datasets within each ARDS model. In the AP group, 34 shared proteins were identified (Figure 5A), primarily enriched in extracellular space and collagen-containing extracellular matrix components. These proteins were functionally linked to serine-type endopeptidase inhibitor activity and acute-phase and innate immune responses. Among these, 21 proteins were consistently upregulated and 3 proteins were downregulated in both tissue and plasma EV compartments (Figure 5B).

**Figure 5.**
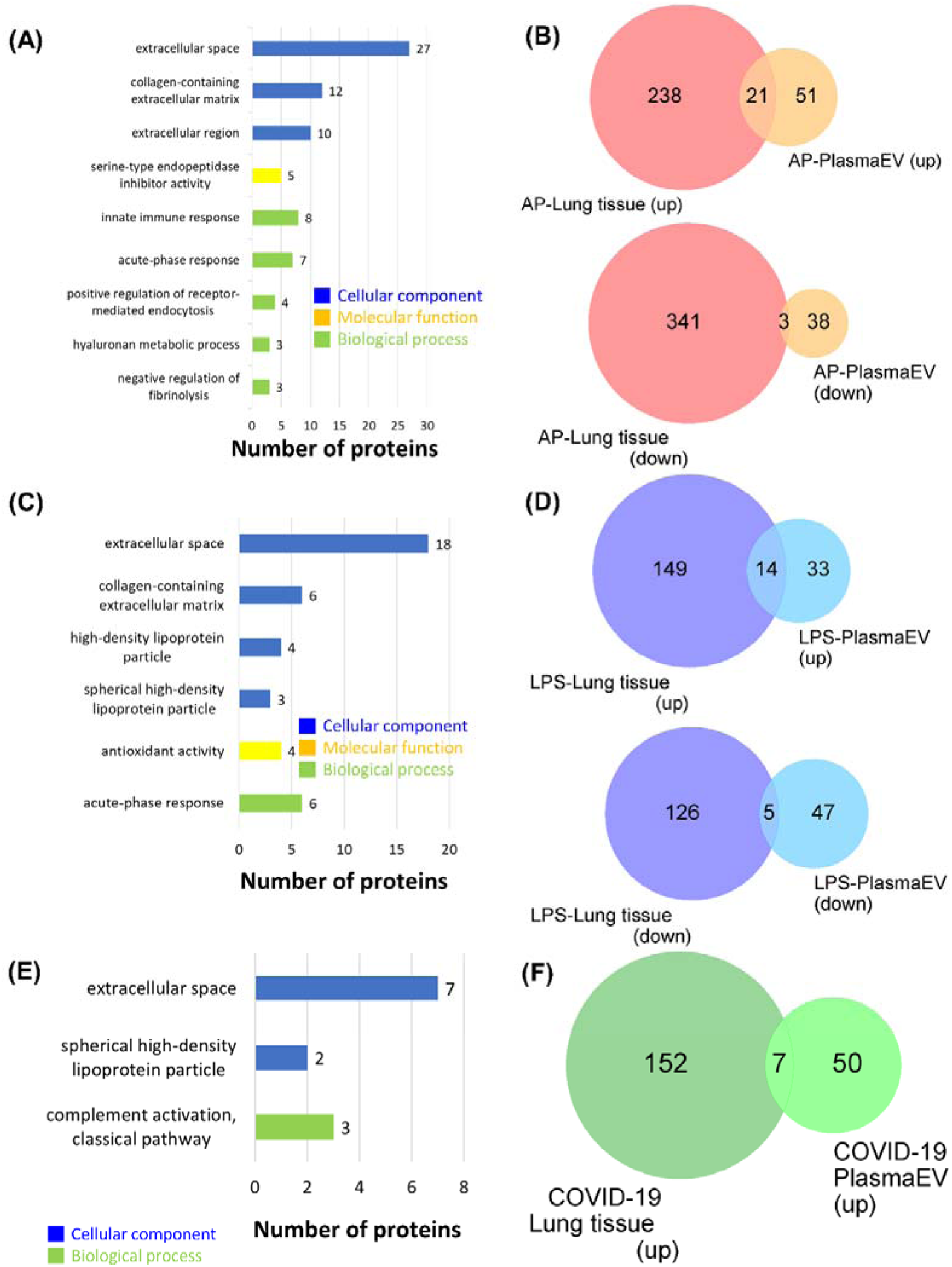
Gene ontology (GO) enrichment analysis tissue-guided correlation between lung and plasma-EV proteomes. The bar plots on the right depict enriched Gene Ontology (GO) terms in cellular component, molecular function, and biological process categories for the overlapping DEPs. (A) Functional classification of differentially abundant proteins in the intersection of lung tissue and plasma-EV samples of the AP group of ARDS mice based on Gene Ontology. (B) Venn diagrams illustrating the overlap among differentially expressed proteins from the intersection of lung tissue and plasma-EV samples with the AP group of ARDS mice, significantly increased proteins (upper), and significantly decreased proteins (lower). (C) Functional classification of differentially abundant proteins in the intersection of lung tissue and plasma-EV samples of the LPS group of ARDS mice based on Gene Ontology. (D) Venn diagrams illustrating the overlap among differentially expressed proteins from the intersection of lung tissue and plasma-EV samples with the LPS group of ARDS mice, significantly increased proteins (upper), and significantly decreased proteins (lower). (E) Functional classification of differentially abundant proteins in the intersection of lung tissue and plasma-EV samples of the COVID-19 group of ARDS mice based on Gene Ontology. (F) Venn diagrams illustrating the overlap among differentially expressed proteins from the intersection of lung tissue and plasma-EV samples with the COVID-19 group of ARDS mice, significantly increased proteins (upper), and significantly decreased proteins (lower).

In the LPS model, 24 shared proteins were detected (Figure 5C), showing enriched extracellular matrix organization and antioxidant-related molecular functions, with the acute-phase response as the dominant biological process. In both sample types, 14 proteins were commonly upregulated and 5 proteins were downregulated (Figure 5D).

The COVID-19-induced model exhibited a smaller overlap, with 8 shared proteins (Figure 5E) associated with extracellular space and high-density lipoprotein particles. Functional annotation highlighted complement activation via the classical pathway. Seven of these proteins were co-upregulated in the lung and EV datasets, whereas no consistently downregulated proteins were observed (Figure 5F).

Collectively, these results demonstrated that tissue-guided proteomic analysis can identify overlapping yet etiology-specific molecular pathways. The AP and LPS models shared signatures of matrix remodeling and inflammatory activation, whereas the COVID-19 model showed complement-related enrichment, underscoring distinct systemic responses reflected in plasma-EV proteomes.

### 3.5 Dysregulation of arachidonic acid metabolism in ARDS models

To identify metabolic pathways commonly reflected in lung tissues and circulating EVs, differential metabolite sets from both sources were intersected within each ARDS model. In the AP group, 606 shared metabolites were identified (Figure 6A) which were significantly enriched in arachidonic acid, galactose, and D-amino acid metabolism pathways (Figure 6B). The LPS model revealed 360 overlapping metabolites (Figure 6C), with enrichment in arachidonic acid, galactose, and fructose–mannose metabolism (Figure 6D). In the COVID-19-induced group, 353 shared metabolites were detected (Figure 6E) which were predominantly associated with arachidonic acid, sphingolipid, and pyruvate metabolism (Figure 6F).

**Figure 6.**
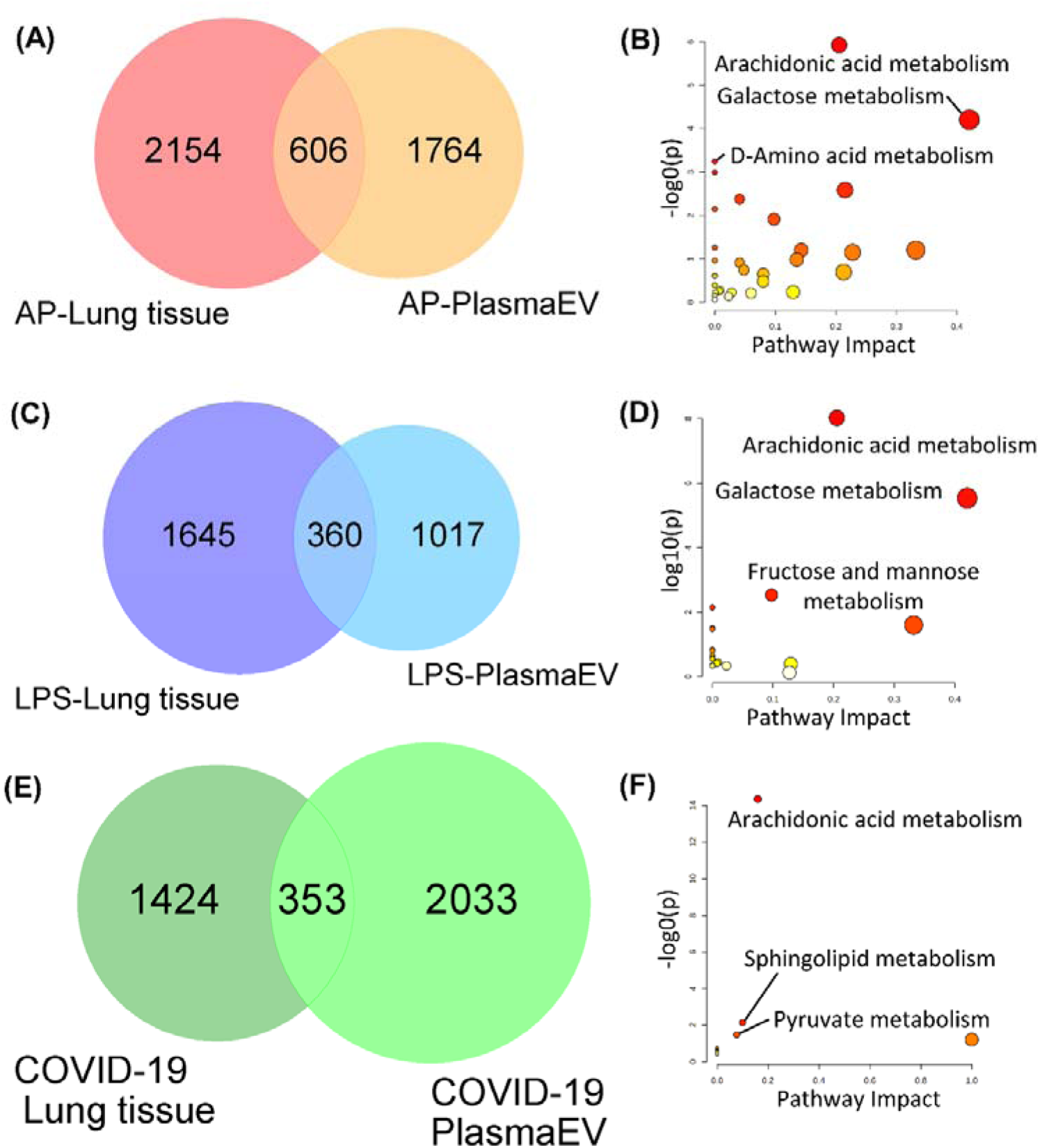
KEGG pathway analysis of tissue-guided correlation between lung and plasma-EV metabolomes. Identified metabolites were analyzed by the “Kyoto Encyclopedia of Genes and Genomes” (KEGG) from pathway analysis. (A) Venn diagram of total DEMs in plasma-EVs and lung tissues in the AP group. (B) The top three significantly enriched pathways were analyzed by pathway analysis of differentially expressed metabolites at the intersection of plasma-EV and lung tissue in the AP group. (C) Venn diagram of total DEMs in plasma-EVs and lung tissues in the LPS group. (D) The top three significantly enriched pathways were analyzed by pathway analysis of differentially expressed metabolites at the intersection of plasma-EV and lung tissue in the LPS group. (E) Venn diagram of total DEMs in plasma-EVs and lung tissues in the COVID-19 group. (F) The top three significantly enriched pathways were analyzed by pathway analysis of differentially expressed metabolites at the intersection of plasma-EV and lung tissue in the COVID-19 group.

Across all three etiologies, arachidonic acid metabolism consistently emerged as a top-ranked pathway, indicating that lipid-mediated inflammatory signaling is a unifying molecular feature of ARDS. The detailed pathway names, raw and transformed *p* values for each comparison are provided in Table S2. Together, these findings highlight that dysregulation of arachidonic acid-related metabolites is a hallmark event reflected in both local lung tissue and systemic EV compartments across diverse ARDS triggers.

### 3.6 Integrated multiomics network analysis of ARDS subtypes

To elucidate the systems-level interactions underlying ARDS, multiomics integration of proteomic and metabolomic data was performed using OmicsNet. In the AP model, the merged network comprising 509 metabolites, 795 proteins, and 2,522 edges was derived from DEPs and DEMs in lung tissues (Figure 7A–C). KEGG-based functional enrichment showed significant activation of glycolysis/gluconeogenesis, arachidonic acid metabolism, and galactose metabolism pathways (*p* < 0.05), indicating enhanced energy turnover and inflammatory lipid remodeling.

**Figure 7.**
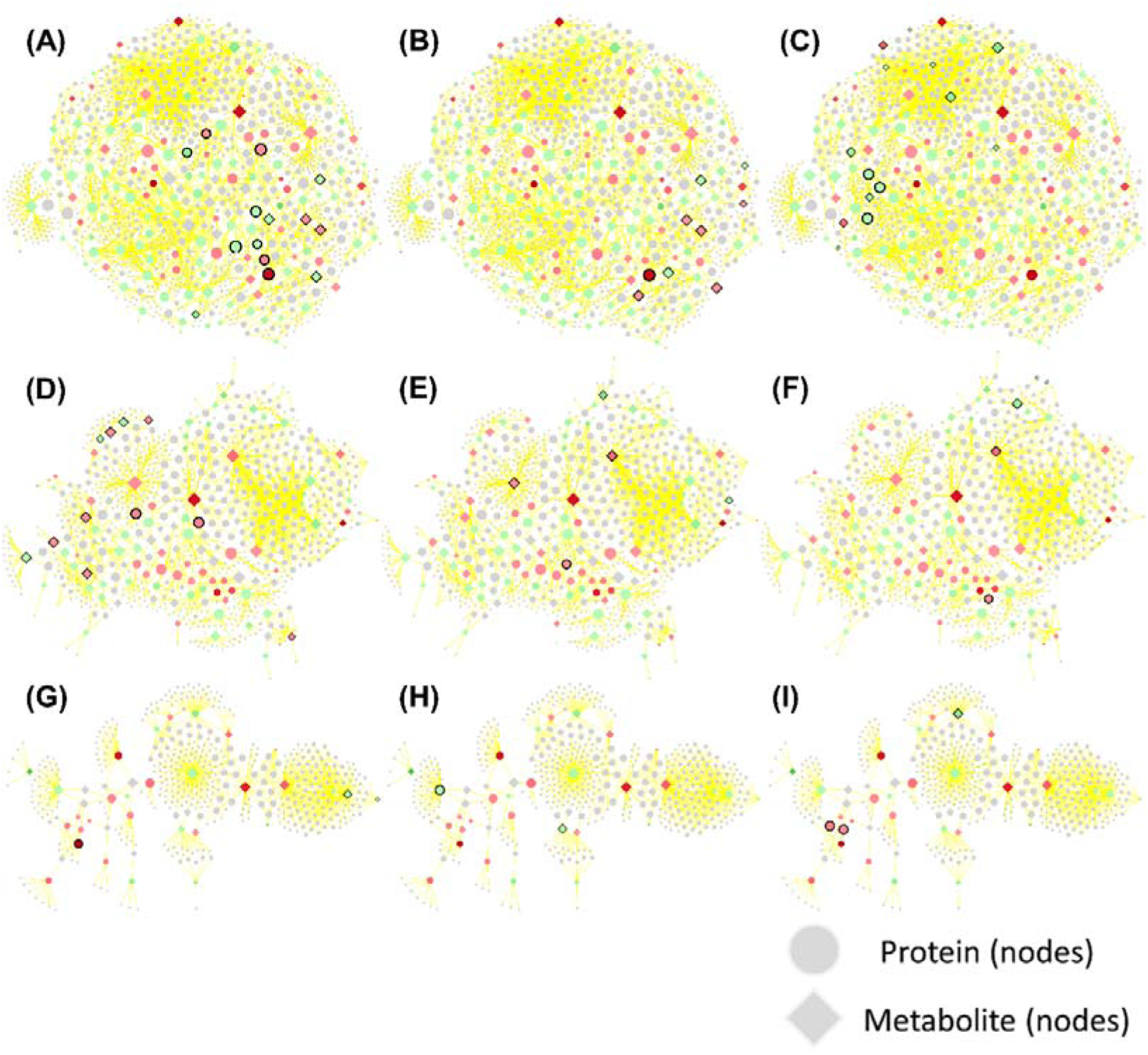
Pilot results of integrated proteomics and metabolomics data revealed enriched pathways for lung tissues. We used an ARDS groups versus a control group as a model to optimize the nitration parameters. Proteins and metabolites are presented as gray dots and diamonds. Symbols with red color were upregulated targets identified in our platform; Symbols with green color were down-regulated targets identified in our system. Symbols with black margins were proteins and metabolites fitted into the respective pathway, which was analyzed from our regulated omics data, in this subnetwork. (A) Top 1 pathway: Glycolysis/Gluconeogenesis in AP group. (B) Second pathway: Amino sugar and nucleotide sugar metabolism in AP group. (C) Third pathway: Arachidonic acid metabolism in AP group. (D)) Top 1 pathway: Platelet activation in LPS group. (E) Second pathway: Regulation of lipolysis in adipocytes in LPS group. (F) Third pathway: ABC transporters in LPS group. (G) Top 1 pathway: Gastric cancer in COVID-19 group. (H) Second pathway: Salmonella infection in COVID-19 group. (I) Third pathway: Vascular smooth muscle contraction in COVID-19 group.

In the LPS model, the integrated network included 244 metabolites, 529 proteins, and 1,411 edges (Figure 7D–F). Enrichment analysis identified platelet activation, regulation of lipolysis in adipocytes, and ABC transporter pathways as top-ranked features, suggesting the involvement of coagulation and lipid-transport processes in sepsis-induced lung injury.

The COVID-19-induced model exhibited a smaller and less connected subnetwork (116 metabolites, 351 proteins, and 667 edges; Figure 7G–I). Functional annotation indicated enrichment in pathways associated with vascular smooth muscle contraction, Salmonella infection, and gastric cancer signaling indicated endothelial and immune dysregulation, which was distinct from bacterial or aspiration models.

Collectively, these integrated networks showed that although each ARDS subtype possessed a unique molecular interaction landscape, these also shared partial convergence in glycolytic and lipid metabolic pathways. Complete pathway lists and joint p-values are provided in Table S3.

### 3.7 Identification of potential plasma EV biomarkers guided by lung tissue profiles

To identify common molecular signatures across ARDS etiologies, intersection analysis of upregulated and downregulated proteins from plasma-EV and lung tissue datasets was performed. Four proteins—haptoglobin (HP), inter-alpha-trypsin inhibitor heavy chain 4 (ITIH4), inter-alpha-trypsin inhibitor heavy chain 3 (ITIH3), and clusterin (CLU)-were consistently upregulated across all ARDS models in both compartments (Figure 8A). Their normalized LFQ intensities showed concordant elevation in lung tissue and plasma EVs (Figure 8C), supporting the robustness of these markers as tissue-guided circulating biomarkers. Only carbonic anhydrase 3 (CA3) exhibited concordant downregulation in the AP and LPS models, whereas no commonly downregulated proteins were shared in the COVID-19 group (Figure 8B). Integrative analysis of metabolomic intersections further identified 77 shared metabolites present in all ARDS models (Figure 8C). These compounds comprised six distinct molecular formulas: C□H□□O□ (18 metabolites), C□□H□□O□ (38), C□□H□□O□ (2), C□□H□□O□ (13), C□□H□□O□ (1), and C□H□□ (5). Box-plot comparison of normalized abundances (Figure 8D) revealed that metabolites with formulas C□H□□O□ and C□□H□□O□, corresponding to oxo-carboxylic acids and eicosanoid-related lipid mediators were consistently downregulated in both lung and EV samples across all models, whereas the remaining four chemical classes displayed discordant regulation patterns.

**Figure 8.**
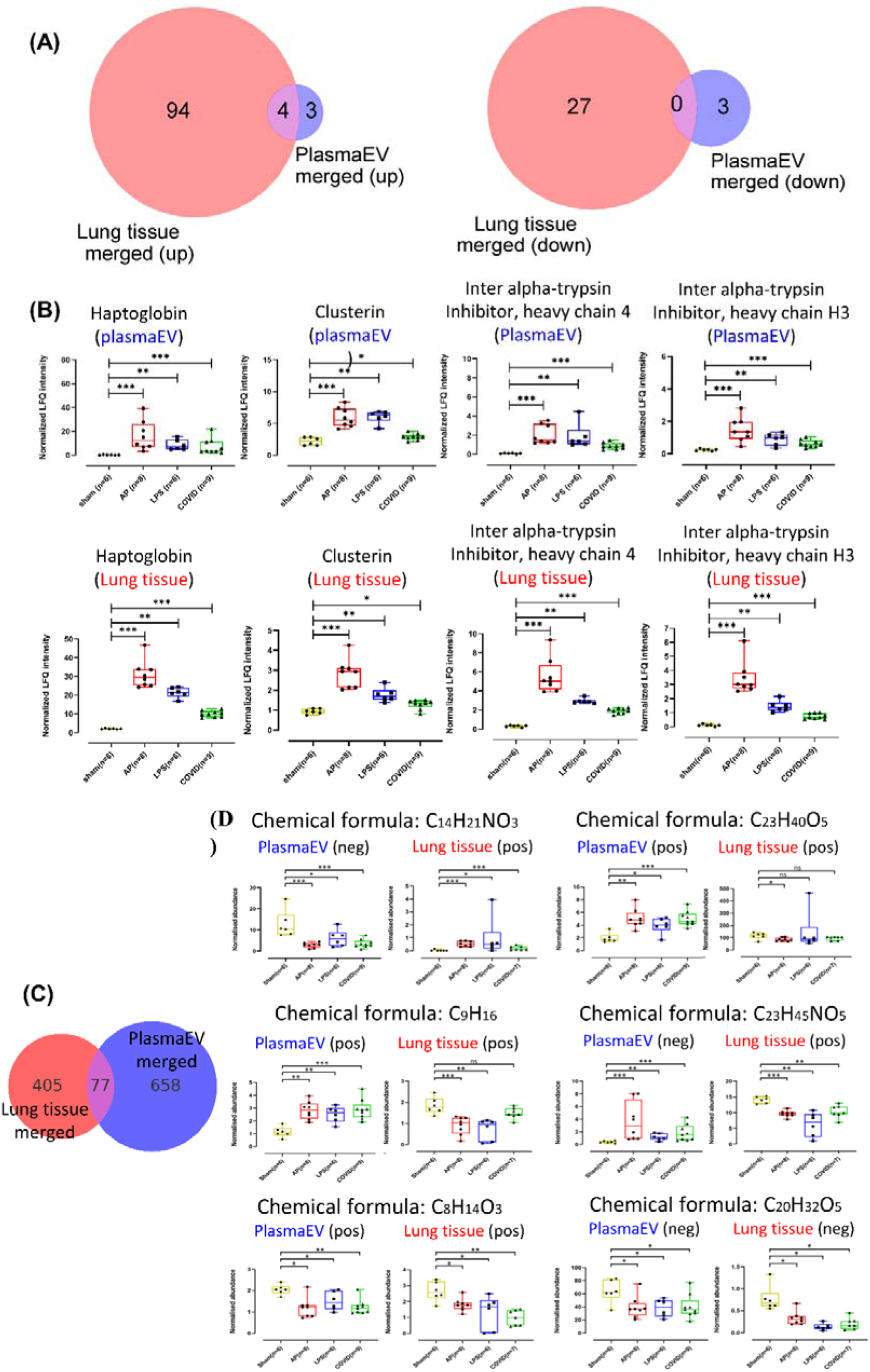
Identification of potential plasma-EV biomarkers of proteins and metabolites guided by lung tissue profiles. The Mann-Whitney test was conducted after ruling out the outliers. The p values were presented with asterisks (ns: P > 0.05, *: P ≤ 0.05, **: P ≤ 0.01, ***: P ≤ 0.001, ****: P ≤ 0.0001). (A)Venn diagrams showing overlapping DEPs between lung tissues and EVs in those ARDS groups. Venn diagrams show all up-regulated proteins with both in plasma-EV and lung tissue samples (left). (B) Four plasma-EV protein biomarkers are showed with the box plot from both plasma-EV and lung tissue samples. (C) Venn diagrams showing overlapping DEMs between lung tissues and EVs in those ARDS groups. (D) Normalised abundance data with the chemical formula are showed with the box plot from both plasma-EV and lung tissue samples.

These results highlight convergent molecular readouts between lung tissue and circulating EVs and pinpoint HP, ITIH4, ITIH3, CLU, and selected eicosanoid-derived metabolites as promising plasma EV biomarkers reflective of lung-specific pathology in ARDS. Full metabolite information is provided in Table S4.

## 4. DISCUSSION

We would first want to focus on the integrated proteomics data from plasma EV and lung tissue samples in ARDS mice models. We found that 4 proteins were up-regulated in the three types of ARDS models in both plasma EV and lung tissue samples. Hp, which is primarily synthesized by the liver, plays a key role in capturing free hemoglobin released by destroyed red blood cells in the body, thereby mitigating its harmful effects. Hp levels decrease in the presence of large amounts of free heme and are therefore a hallmark of hemolysis [23]. Other studies have shown that changes in the concentration of hemoglobin and hemopexin protein in the human body caused by hemolysis are also associated with a worse prognosis of sepsis and ARDS [24]. Gong et al. also found that some differentially expressed proteins like SAA1, SAA2, and Hp are all up-regulated in both serum and lung tissue samples [15]. The inter-alpha-trypsin inhibitor family not only protects the extracellular matrix through protease inhibition but also interacts with matrix molecules such as hyaluronan, suppresses complement activation, and influences cell signaling pathways [25]. Although research on inter-alpha-trypsin inhibitor family protein in ARDS is limited, several studies have demonstrated that inter-alpha-trypsin inhibitor family proteins promote the repair of injured bronchial epithelial cells and mitigate lung injury by binding to vitronectin [26], and suppressing complement activation [27]. CLUs are ubiquitous in mammalian tissues and body fluids and exist in a variety of forms and has many functions in different organs [28]. There are no studies investigating CLU in ARDS. Evidence suggests that dysregulation of the concentration of CLU and transforming growth factor beta 1 is associated with the pulmonary fibrosis [29]. Pulmonary fibrosis is one of the symptoms in ARDS [30].

We next want to focus the discussion on the integrated metabolomics data from plasma EV and lung tissue samples in ARDS mice models. This study results showed that arachidonic acid metabolism pathway to be significant in all three types of ARDS mice models plasma-EV and lung tissue samples. Several studies have suggested that arachidonic acid plays an important role in some diseases including cardiovascular biology [31], carcinogenesis [32], and many inflammatory diseases [33]. Chang et al. collected human plasma from sepsis-ARDS patients and healthy controls to analyze the metabolic alterations between these two groups. The result showed that arachidonic acid metabolism was one of the most significant pathways associated with sepsis-induced ARDS [34]. These findings are similar to the current study data.

Previous studies have reported that suppressing COX-2, an enzyme that plays a key role in the arachidonic acid metabolic pathway, attenuates LPS-induced ALI [35]. However, many studies found that arachidonic acid metabolism-associated metabolites were up-regulated in most of the ARDS mice models [36, 37]. However, our study results data with arachidonic acid metabolism-associated metabolites were contrary to the previous findings. Given that several metabolites are regulated through the same metabolic pathway, it cannot be established that the entire metabolic pathway is upregulated.

Lastly, the multiomics results showed different insights into the three ARDS mice models. In the AP group of the ARDS model, glycolysis was identified to be the most significant pathway. According to the study by Yan et al., elevated lactate levels might reflect enhanced anaerobic glucose metabolism in the community-acquired pneumonia group with accompanying ARDS [38]. Lactate plays a central role in this metabolic pathway. Its accumulation typically occurs when cellular energy and oxygen demands surpass the availability leading to an upregulation of glycolytic activity. Numerous papers have cited the importance of lactate in acute lung disease [39, 40]. Subsequently, in the LPS group, platelet aggregation was identified as the most significant pathway in the multiomics analysis. Many studies have mentioned that platelets are involved in the pathogenesis of ARDS. Studies have found that the levels of platelet-specific α-granules in bronchoalveolar lavage fluid samples in ARDS patients are elevated [41] suggesting that increased platelet activity leads to platelet aggregation. Several studies have reported that, in comparison to critically ill patients without acute lung injury, those with acute respiratory failure exhibited enhanced platelet activation and impaired hemostatic function suggesting altered platelet activity in individuals with ARDS [42]. In the context of systemic conditions that lead to ARDS such as sepsis or cardiopulmonary bypass, some researchers have investigated a pig model of cardiopulmonary bypass and observed that attenuation of platelet aggregation and activity can mitigate lung tissue injury [43]. Thus, platelet aggregation is associated with ARDS. In ARDS model, the COVID-19 group had lesser available multiomics data compared with the other two groups which was likely due to the exhibition of fewer differentially expressed proteins. Despite this, our multiomics analysis identified vascular smooth muscle contraction as the most significantly enriched pathway in the COVID-19 group. Although this pathway is not directly associated with ARDS, some studies have suggested potential links between vascular smooth muscle contraction and the pathophysiology of ARDS [44].

The study has several limitations. First, there were certain challenges in the adduct assignment for compound identification in the metabolomics dataset. In accordance with previously published studies [44–46], only common adduct forms—[M+H]□, [M+Na]□, [M+K]□, and [M+2H]²□ in positive ion mode, and [M–H]□ in negative ion mode—were included to ensure consistency. However, we observed that a single m/z value could correspond to multiple entries in the HMDB, and conversely, one HMDB compound could match several m/z values. These ambiguities reflect inherent limitations in mass-based identification as isomeric or isobaric metabolites cannot be reliably distinguished without additional structural confirmation. Future studies incorporating tandem MS/MS spectral libraries or authentic standards will be necessary to resolve these issues.

## 5. CONCLUSIONS

This study established a tissue-guided multiomics framework that bridges local lung pathology with systemic extracellular vesicle (EV) profiles across distinct ARDS etiologies. By integrating proteomic and metabolomic datasets, we identified four conserved proteins (HP, ITIH3, ITIH4, and CLU) and arachidonic acid–related metabolites as robust circulating indicators of pulmonary injury. These findings demonstrate that plasma EVs not only reflect lung-specific molecular signatures but also delineate etiology-dependent pathogenic networks, including glycolytic activation, platelet aggregation, and vascular smooth muscle dysregulation. Collectively, our work positions tissue-informed EV profiling as a powerful approach to uncover mechanistic and diagnostic biomarkers in ARDS. This framework provides a foundation for developing clinically translatable EV-based assays that could enable early detection, molecular subtyping, and precision monitoring of acute lung injury.

## List of abbreviations

ARDS: acute respiratory distress syndrome
ALI: acute lung injury
EV: extracellular vesicle
EVs: extracellular vesicles
AP: aspiration pneumonia
LPS: lipopolysaccharide
COVID-19: coronavirus disease 2019
HP: haptoglobin
ITIH3: inter-alpha-trypsin inhibitor heavy chain 3
ITIH4: inter-alpha-trypsin inhibitor heavy chain 4
CLU: clusterin
CA3: carbonic anhydrase 3
sPLS-DA: sparse partial least squares discriminant analysis
DEP: differentially expressed protein
DEPs: differentially expressed proteins
DEM: differentially expressed metabolite
DEMs: differentially expressed metabolites
DEF: differentially expressed feature
GO: gene ontology
KEGG: Kyoto Encyclopedia of Genes and Genomes
MS: mass spectrometry
LC–MS/MS: liquid chromatography–tandem mass spectrometry
NTA: nanoparticle tracking analysis
TEM: transmission electron microscopy
PVDF: polyvinylidene difluoride
BCA: bicinchoninic acid
SDS–PAGE: sodium dodecyl sulfate polyacrylamide gel electrophoresis
SPE: solid-phase extraction
HMDB: Human Metabolome Database
RIPA: radioimmunoprecipitation assay
PBS: phosphate-buffered saline
FWHM: full width at half maximum
FC: fold change
COX-2: cyclooxygenase-2.

## Ethics approval

The Institutional Animal Use and Care Committee of Taipei Medical University approved all animal studies (LAC-2022-0427).

## Consent for publication

Not applicable

## Data availability

The datasets generated and/or analyzed during the current study are available from the corresponding author on reasonable request.

## Author contributions

Conceptualization: Y-CH, C-JH, I-LT, Y-JF, Y-WL; Methodology: P-TC, C-YC, C-LC, C-HL, C-WS, C-FL; Investigation: P-TC, C-YC; Formal analysis: C-FL, Y-JF; Data curation: P-TC, C-YC, C-FL; Resources: Y-JF, Y-WL, C-JH; Funding acquisition: Y-CH, C-JH, I-LT, Y-JF, Y-WL; Supervision: Y-CH, C-JH, I-LT; Writing – original draft: P-TC, C-YC; Writing – review & editing: Y-CH, C-JH, I-LT.

## Acknowledgements

The authors are grateful for the technical support provided by the TMU Core Facility, and also the mass spectrometry technical research services from Consortia of Key Technologies and Instrumentation Center, National Taiwan University.

