## Supplemental Figure 1-10, Table 1-4 for "Tissue-guided multi-omics profiling identifies extracellular vesicle biomarkers indicative of lung pathology in acute respiratory distress syndrome"

**Supplementary data**

1. **SUPPLEMENTARY INFORMATION: MATERIALS AND METHOD**

**Material for Proteomics**

The materials for proteomics included SEC columns and the SmartSEC EV Isolation System (System Biosciences) to isolate extracellular vesicles from plasma. Protein quantification was performed using the Pierce™ BCA Kit (Thermo Fisher). Amicon® Ultra-15 filter units (Merck Millipore) were used for concentration, and Protein LoBind tubes (Eppendorf) minimized protein loss. RIPA buffer (Merck Millipore) facilitated lysis, while urea (Nihon Shiyaku), DTT, IAA, ammonium bicarbonate, and formic acid (Sigma-Aldrich) aided in sample preparation. Solvents such as acetone (Honeywell), acetonitrile (J.T. Baker), and methanol (Macron Fine Chemicals) were employed. PBS (VWR), trypsin (Promega), and C18 columns (Waters) completed the workflow.

**LC–MS/MS parameter settings for proteomics**

Tryptic peptides were analyzed using an Ultimate 3000 nanoLC system coupled with an Orbitrap Fusion Lumos mass spectrometer. Peptides were separated on a C18 PepMap column with a 40-minute gradient for plasma and 50-minute for lung tissue, using 0.1% formic acid in water (A) and 100% acetonitrile with 0.1% formic acid (B) at 300 nL/min. MS1 scans were acquired at 120,000 resolution (m/z 200) with a 5e5 AGC target and 50 ms injection time. The top intense ions were fragmented by HCD at 15,000 resolution with a 1.4 Da isolation window and 32 NCE, followed by dynamic exclusion for 60 seconds.

**Material for Metabolomics**

The materials used for the experiments were sourced from various reputable suppliers. Protein LoBind tubes, which minimize protein loss during storage and handling, were sourced from Eppendorf (Hamburg, Germany). Acetone, an essential reagent for protein precipitation, was purchased from Honeywell (Charlotte, NC, USA) under the Burdick & Jackson® brand (AH010-4A). Methanol, specifically AR® Methyl Alcohol, Anhydrous, was sourced from Macron Fine Chemicals™ (Center Valley, PA, USA)

**LC–MS/MS parameter settings for metabolomics**

The LC-MS/MS analysis was carried out using SYNAPT XS (Waters, US). Each sample was injected into an Agilent Poroshell 120 EC-C18 column (1.9µM, 2.1*100 mm) at 40°C.The mass spectrometry settings include an MS1 scanning mass range of 50 Da to 1200 Da, with a capillary voltage of 2 kV and a source temperature of 150°C. The sampling cone is set to 40, and the desolvation temperature is 400°C. The cone gas flow is 50 L/hr. The survey scan starts at a mass of 50 Da and ends at 1000 Da, with an intensity threshold of 10,000 and the detection of up to 10 components.

2. **SUPPLEMENTARY INFORMATION: FIGURES**

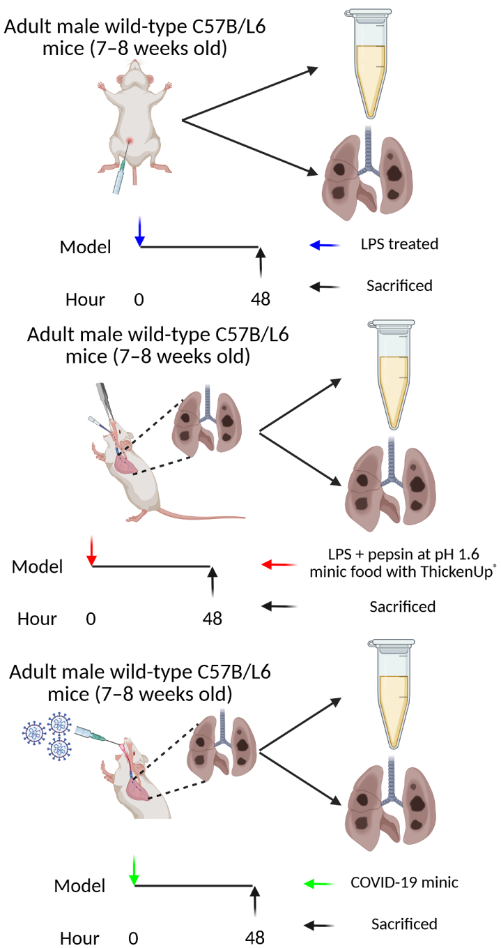

**B**

**A**

**C**

**Supplementary Figure 1. Flow chart of an animal model.**

A. Lipopolysaccharide (LPS group) ARDS model mice were treated. B. Aspiration Pneumonia (AP group) ARDS model mice treated. C. Coronavirus disease 2019 (COVID-19) ARDS model mice treated.

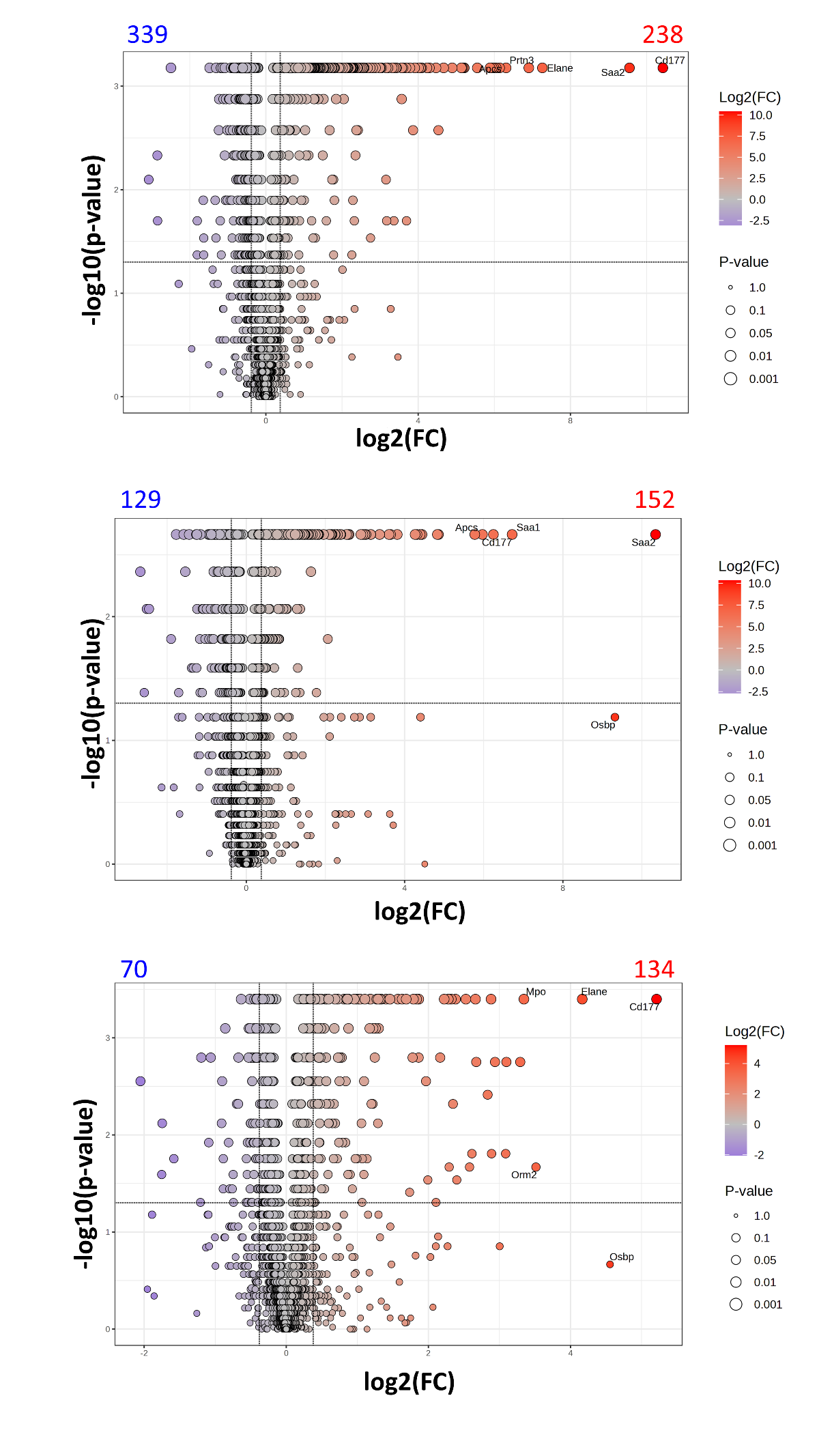

**C**

**B**

**A**

**Supplementary Figure 2.** **Quantitative proteome analysis of ARDS group and sham group from lung tissue samples.**

Proteins from proteomics data with adjusted p-value < 0.05 and fold change > 1.3 were colored. Those numbers of the up-regulation protein were labeled in red. Those numbers of the down-regulation protein were labeled in blue. A. Volcano plot showing the differentially expressed proteins between the AP group and the sham group from lung tissue samples. B. Volcano plot showing the differentially expressed proteins between the LPS group and the sham group from lung tissue samples. C. Volcano plot showing the differentially expressed proteins between the COVID-19 group and sham group from lung tissue samples.

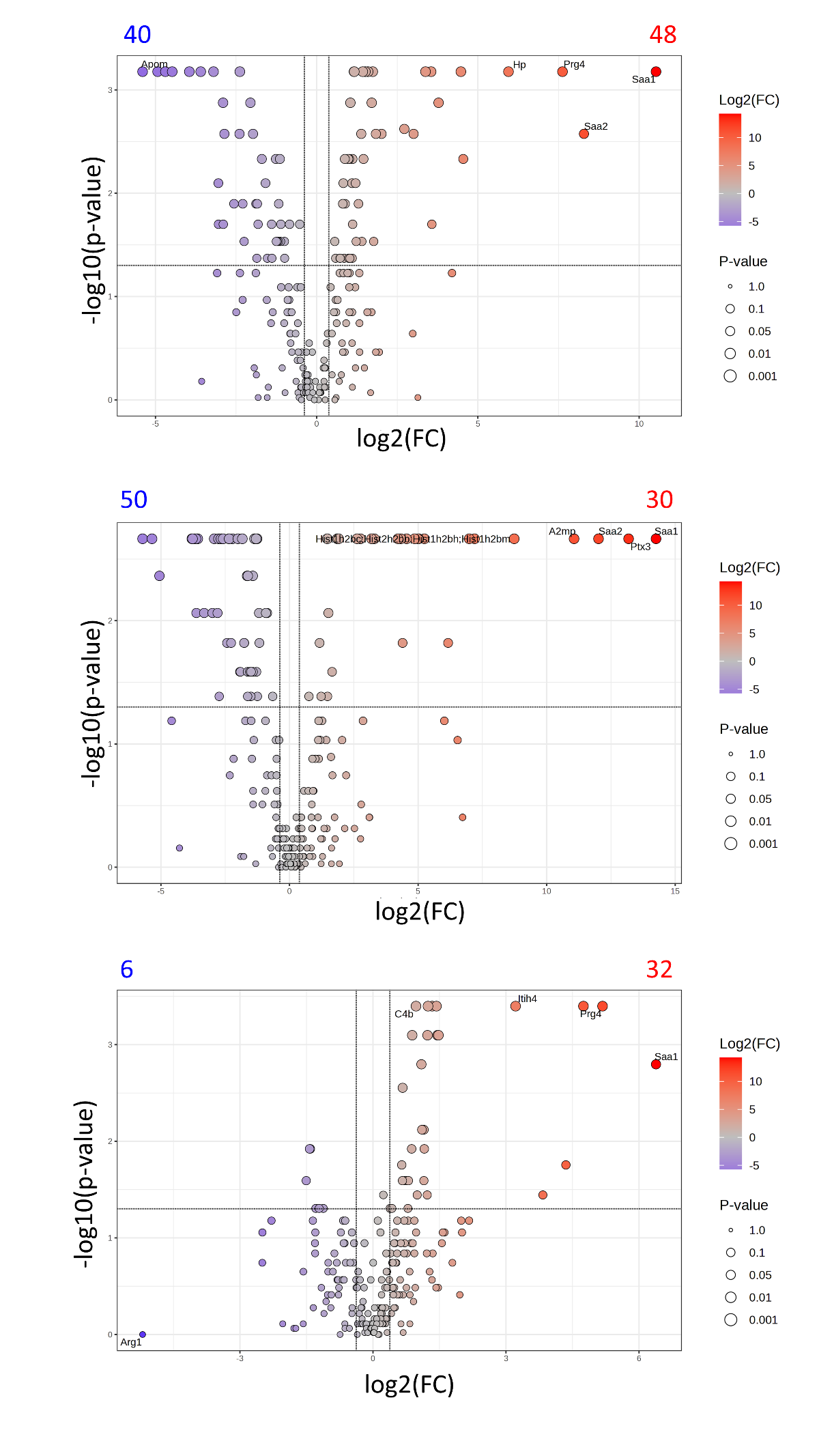

**A**

**B**

**C**

**Supplementary Figure 3**. **Quantitative proteome analysis of ARDS group and sham group from plasma-EV samples.**

Proteins from proteomics data with adjusted p-value < 0.05 and fold change > 1.3 were colored. Those numbers of the up-regulation protein were labeled in red. Those numbers of the down-regulation protein were labeled in blue. A. Volcano plot showing the differentially expressed proteins between the AP group and the sham group from plasma-EV samples. B. Volcano plot showing the differentially expressed proteins between the LPS group and the sham group from plasma-EV samples. C. Volcano plot showing the differentially expressed proteins between the COVID group and sham group from plasma-EV samples.

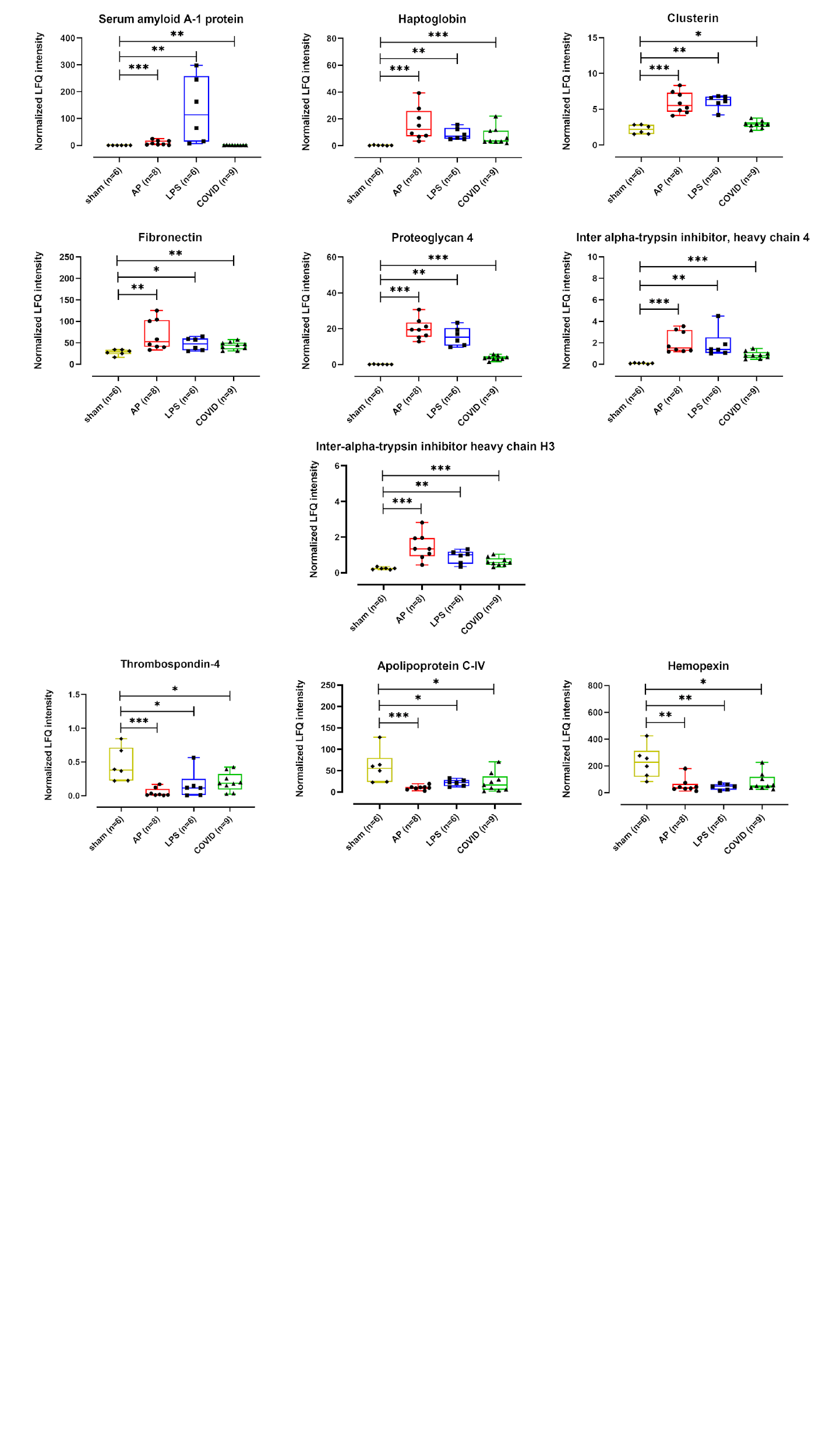

**Supplementary Figure 4**. **Box plot analysis comparing the LFQ intensity of 10 plasma-EV DEPs, which overlapped in the three ARDS models.**

The Mann-Whitney test was conducted after ruling out the outliers. The p values were presented with asterisks (ns: P > 0.05, *: P ≤ 0.05, **: P ≤ 0.01, ***: P ≤ 0.001, ****: P ≤ 0.0001)

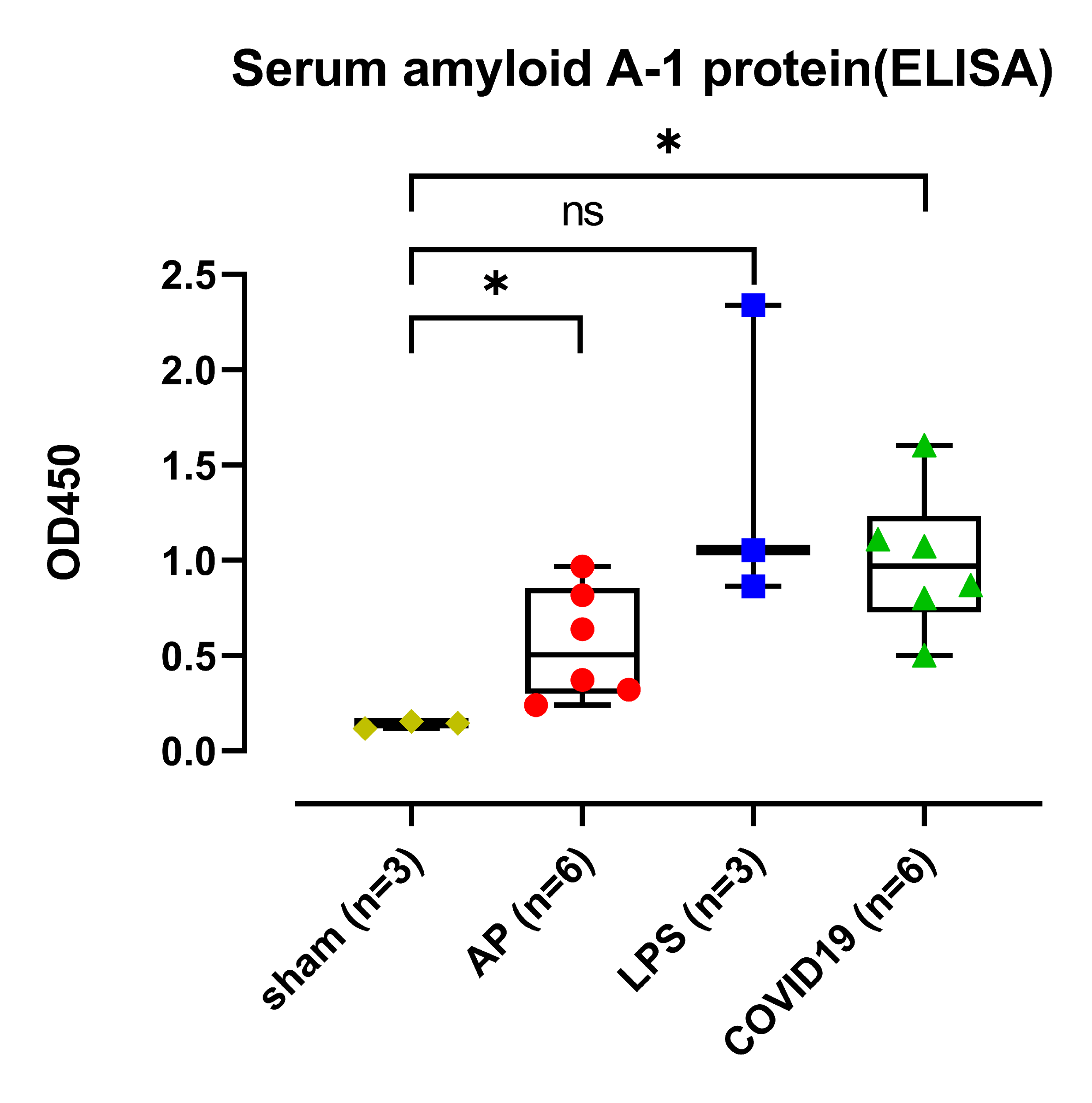

**Supplementary Figure 5**. **ELISA was used to measure the protein expression levels of mouse Serum Amyloid A1 (SAA1) protein.**

Asterisks indicate significant differences between the groups. The Mann-Whitney test was conducted after ruling out the outliers. The *p* values were presented with asterisks (ns: P > 0.05, *: P ≤ 0.05, **: P ≤ 0.01, ***: P ≤ 0.001, ****: P ≤ 0.0001), ELISA, enzyme-linked immunosorbent assay.

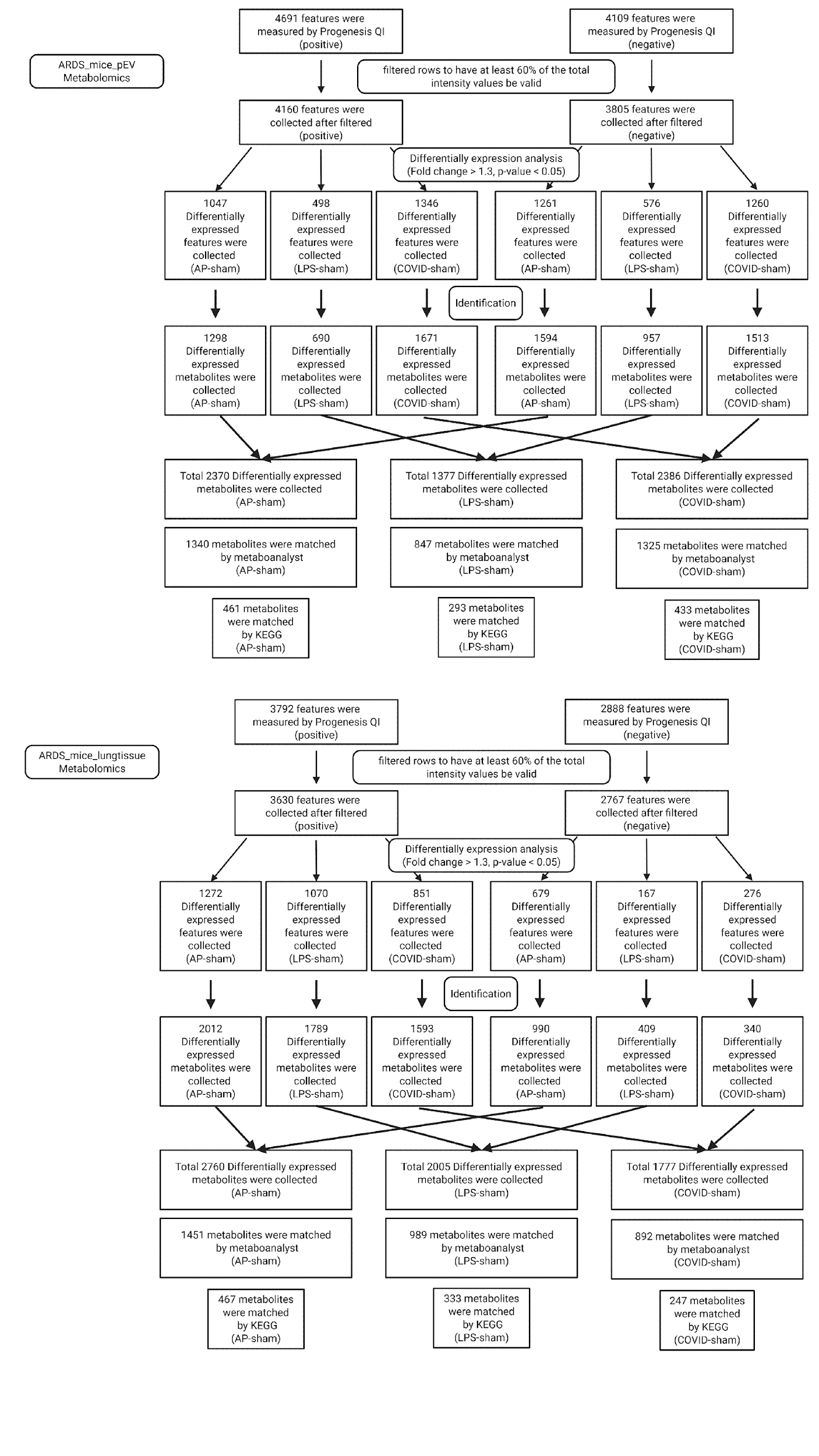

**A**

**B**

**Supplementary Figure 6. Metabolomics data dealing with the ARDS model mice from plasma-EV and lung tissue samples.**

A. Flowchart of features screening and numbers in metabolomics data from the plasma-EV samples of the ARDS model mice. B. Flowchart of features screening and numbers in metabolomics data from the lung tissue samples of the ARDS model mice.

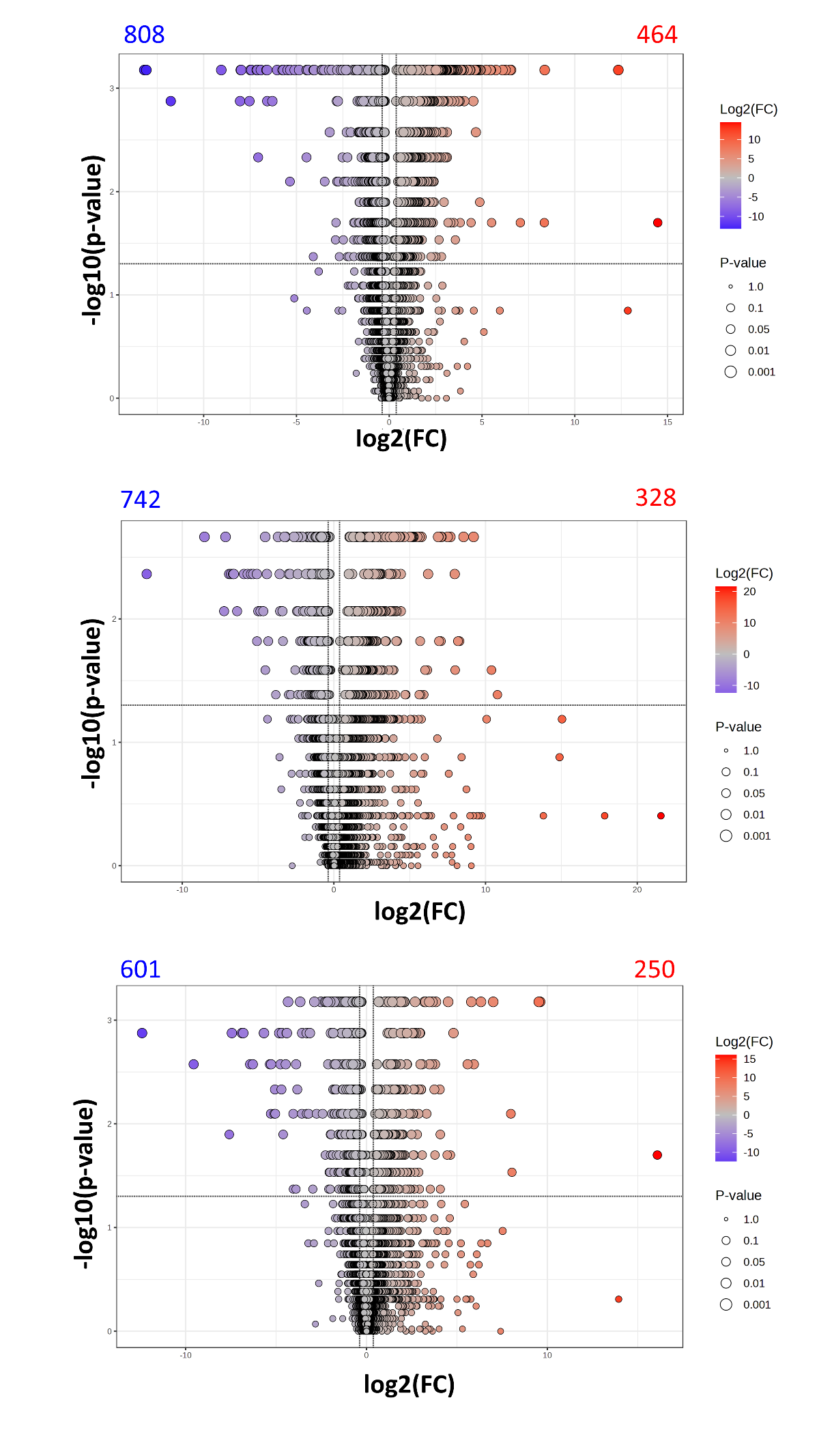

**A**

**B**

**C**

**Supplementary Figure 7. Differentially expressed features of ARDS group and sham group from lung tissue samples in positive mode.**

Features from metabolomics data with adjusted *p* value < 0.05 and fold change > 1.3 were colored. Those numbers of the up-regulation feature were labeled in red. Those numbers of the down-regulation feature were labeled in blue. A. Volcano plot showing the differentially expressed features between the AP group and the sham group from lung tissue samples in positive mode. B. Volcano plot showing the differentially expressed features between the LPS group and the sham group samples’ lung tissue in positive mode. C. Volcano plot showing the differentially expressed features between the COVID-19 group and sham group samples’ lung tissue in positive mode.

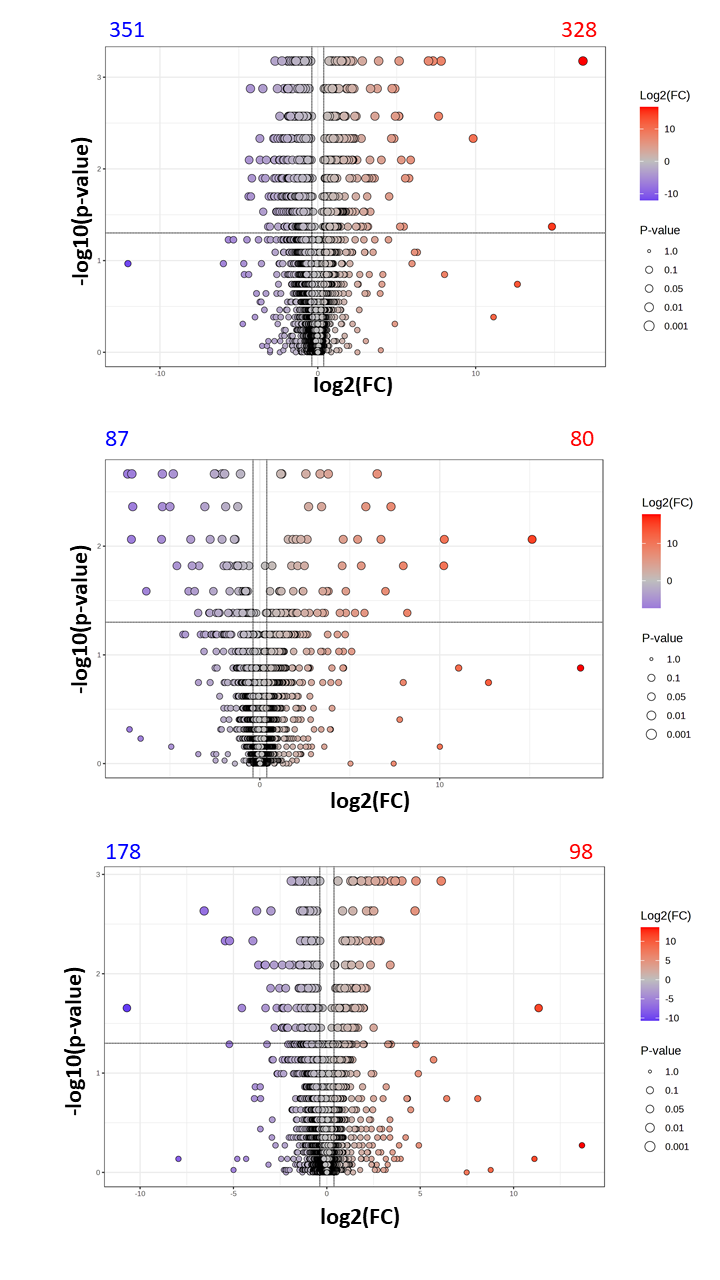

**A**

**B**

**C**

**Supplementary Figure 8. Differentially expressed features of the ARDS group and the sham group from lung tissue samples in negative mode.**

Features from metabolomics data with adjusted *p* value < 0.05 and fold change > 1.3 were colored. Those numbers of the up-regulation feature were labeled in red. Those numbers of the down-regulation feature were labeled in blue. A. Volcano plot showing the differentially expressed features between the AP group and the sham group from lung tissue samples in negative mode. B. Volcano plot showing the differentially expressed features between the LPS group and the sham group from lung tissue samples in negative mode. C. Volcano plot showing the differentially expressed features between the COVID-19 group and sham group from lung tissue samples in negative mode.

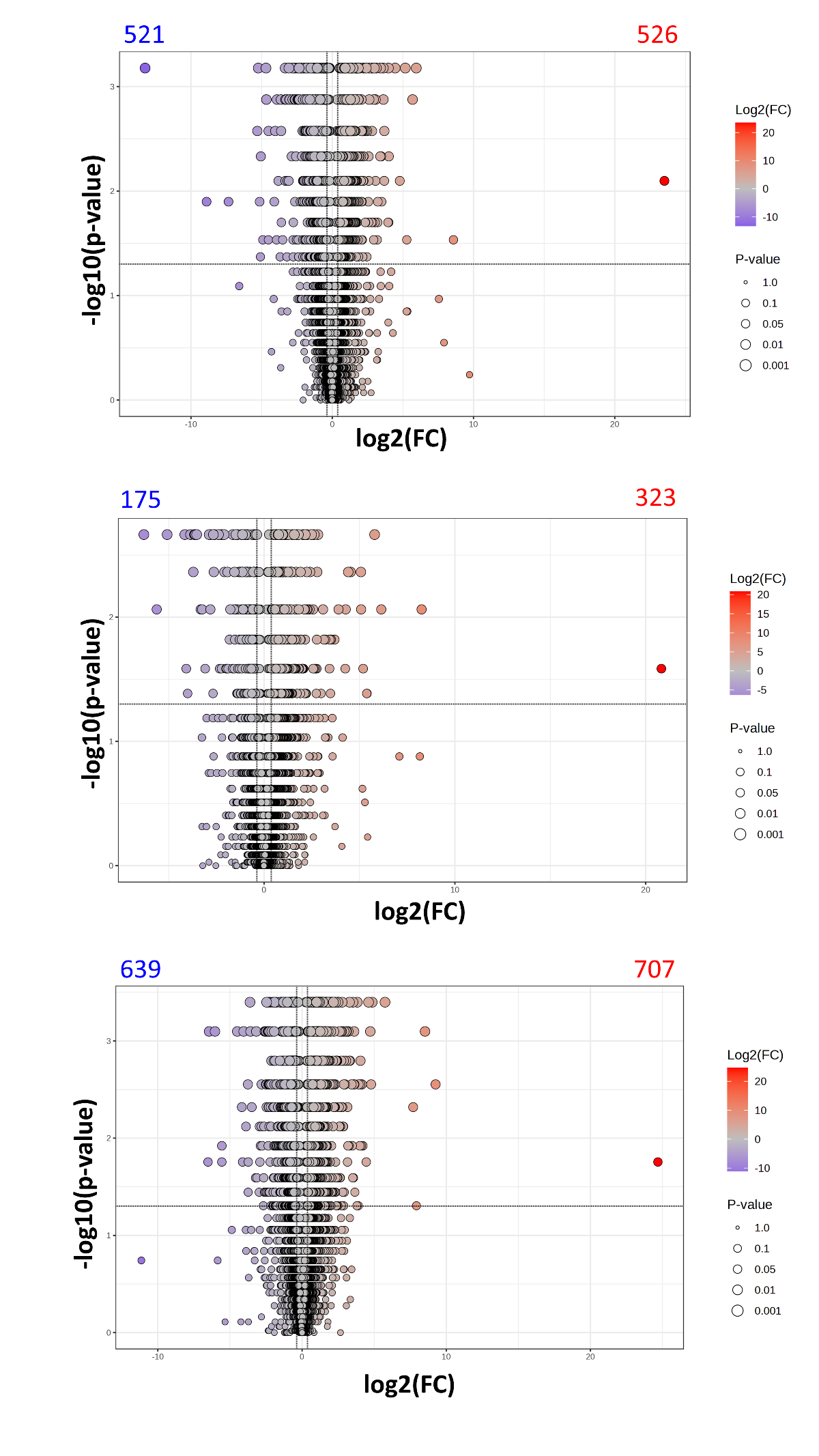

**A**

**B**

**C**

**Supplementary Figure 9. Differentially expressed features of ARDS group and sham group from plasma-EV samples in positive mode.**

Features from metabolomics data with adjusted *p* value < 0.05 and fold change > 1.3 were colored. Those numbers of the up-regulation feature were labeled in red. Those numbers of the down-regulation feature were labeled in blue. A. Volcano plot showing the differentially expressed features between the AP group and the sham group from plasma-EV samples in positive mode. B. Volcano plot showing the differentially expressed features between the LPS group and the sham group from plasma-EV samples in positive mode. C. Volcano plot showing the differentially expressed features between the COVID-19 group and sham group from plasma-EV samples in positive mode.

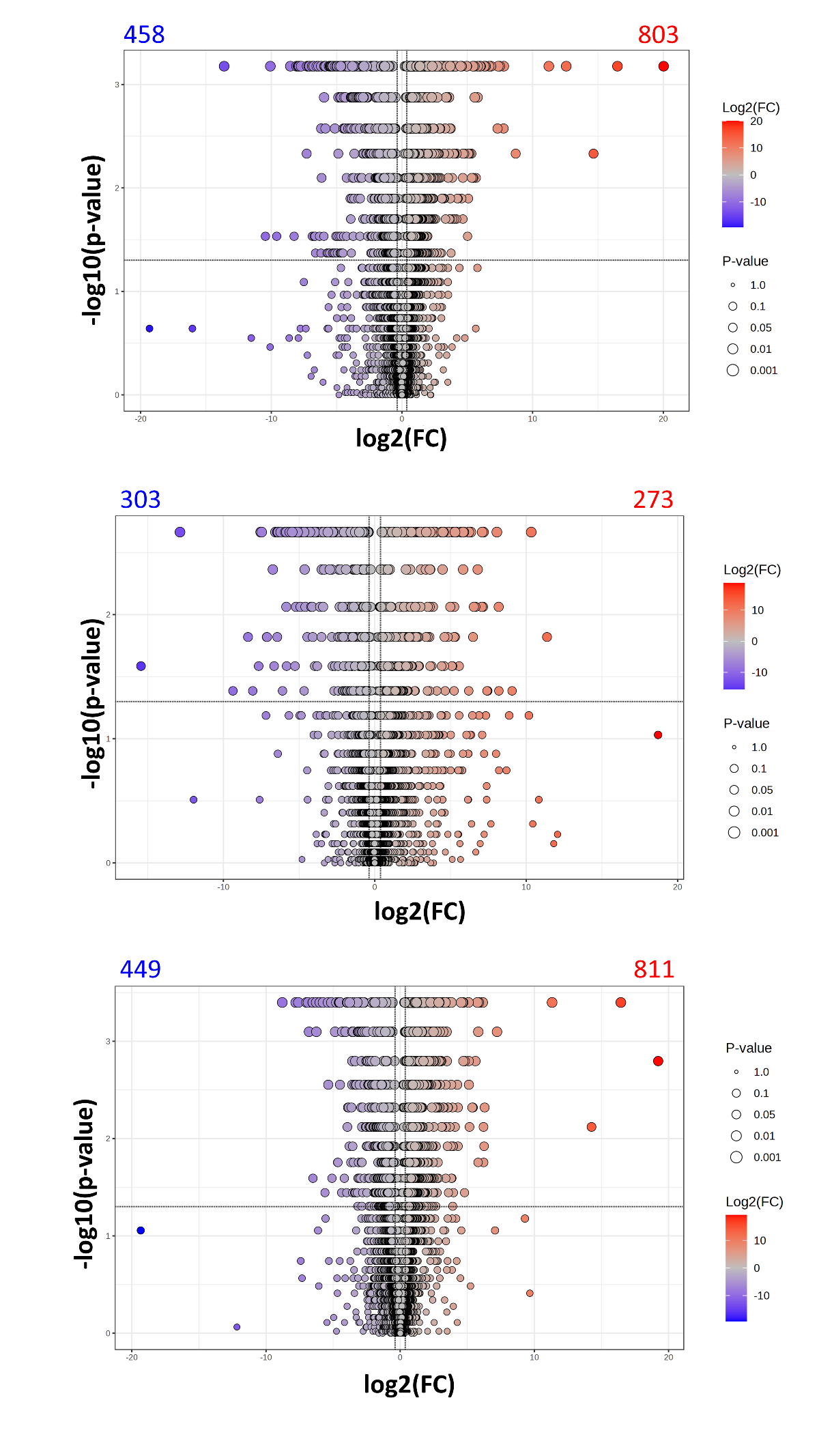

**A**

**B**

**C**

**Supplementary Figure 10. Differentially expressed features of ARDS group and sham group from plasma-EV samples in negative mode.**

Features from metabolomics data with adjusted *p* value < 0.05 and fold change > 1.3 were colored. Those numbers of the up-regulation feature were labeled in red. Those numbers of the down-regulation feature were labeled in blue. A. Volcano plot showing the differentially expressed features between the AP group and the sham group from plasma-EV samples in negative mode. B. Volcano plot showing the differentially expressed features between the LPS group and the sham group from plasma-EV samples in negative mode. C. Volcano plot showing the differentially expressed features between the COVID-19 group and sham group from plasma-EV samples in negative mode.

**Supplementary Table 1. A list of 125 proteins was found by intersection analysis, which is from lung tissue samples, and the proteins that were expressed in both plasma-EV and lung tissue samples are in bold.**

| **UniProt ID** | **UniProt accession** | **Regulation** | **Protein name** |
| --- | --- | --- | --- |
| CD177_MOUSE | Q8R2S8 | UP | CD177 antigen |
| ELNE_MOUSE | Q3UP87 | UP | Neutrophil elastase |
| A1AG2_MOUSE | P07361 | UP | Alpha-1-acid glycoprotein 2 |
| PERM_MOUSE | P11247 | UP | Myeloperoxidase |
| IFIT3_MOUSE | Q64345 | UP | Interferon-induced protein with tetratricopeptide repeats 3 |
| PRTN3_MOUSE | Q61096 | UP | Myeloblastin |
| CY24B_MOUSE | Q61093 | UP | Cytochrome b-245 heavy chain |
| SAA2_MOUSE | P05367 | UP | Serum amyloid A-2 protein |
| NGAL_MOUSE | P11672 | UP | Neutrophil gelatinase-associated lipocalin |
| SAMP_MOUSE | P12246 | UP | Serum amyloid P-component |
| **ITIH3_MOUSE** | **Q61704** | **UP** | **Inter-alpha-trypsin inhibitor heavy chain H3** |
| CATG_MOUSE | P28293 | UP | Cathepsin G |
| MMP9_MOUSE | P41245 | UP | Matrix metalloproteinase-9 |
| **ITIH4_MOUSE** | **A6X935** | **UP** | **Inter alpha-trypsin inhibitor, heavy chain 4** |
| TPSN_MOUSE | Q9R233 | UP | Tapasin |
| ITAM_MOUSE | P05555 | UP | Integrin alpha-M |
| S10A9_MOUSE | P31725 | UP | Protein S100-A9 |
| **HPT_MOUSE** | **Q61646** | **UP** | **Haptoglobin** |
| LG3BP_MOUSE | Q07797 | UP | Galectin-3-binding protein |
| S10A8_MOUSE | P27005 | UP | Protein S100-A8 |
| IIGP1_MOUSE | Q9QZ85 | UP | Interferon-inducible GTPase 1 |
| CD14_MOUSE | P10810 | UP | Monocyte differentiation antigen CD14 |
| IRGM3_MOUSE | Q9DCE9 | UP | Immunity-related GTPase family M protein 3 |
| TRFL_MOUSE | P08071 | UP | Lactotransferrin |
| STAT1_MOUSE | P42225 | UP | Signal transducer and activator of transcription 1 |
| NGP_MOUSE | O08692 | UP | Neutrophilic granule protein |
| CH3L1_MOUSE | Q61362 | UP | Chitinase-3-like protein 1 |
| TGTP1_MOUSE | Q62293 | UP | T-cell-specific guanine nucleotide triphosphate-binding protein 1 |
| TGTP2_MOUSE | Q3T9E4 | UP | T-cell-specific guanine nucleotide triphosphate-binding protein 2 |
| ITB2_MOUSE | P11835 | UP | Integrin beta-2 |
| GBP2_MOUSE | Q9Z0E6 | UP | Guanylate-binding protein 2 |
| CAMP_MOUSE | P51437 | UP | Cathelicidin antimicrobial peptide |
| CHIL3_MOUSE | O35744 | UP | Chitinase-like protein 3 |
| A1AG1_MOUSE | Q60590 | UP | Alpha-1-acid glycoprotein 1 |
| HCLS1_MOUSE | P49710 | UP | Hematopoietic lineage cell-specific protein |
| IRGM1_MOUSE | Q60766 | UP | Immunity-related GTPase family M protein 1 |
| SERA_MOUSE | Q61753 | UP | D-3-phosphoglycerate dehydrogenase |
| RAC2_MOUSE | Q05144 | UP | Ras-related C3 botulinum toxin substrate 2 |
| ISG15_MOUSE | Q64339 | UP | Ubiquitin-like protein ISG15 |
| AMBP_MOUSE | Q07456 | UP | Protein AMBP |
| HB2A_MOUSE | P14483 | UP | H-2 class II histocompatibility antigen, A beta chain |
| HB2F_MOUSE | P06346 | UP | H-2 class II histocompatibility antigen, A-F beta chain |
| HB2U_MOUSE | P06344 | UP | H-2 class II histocompatibility antigen, A-U beta chain |
| HB2K_MOUSE | P06343 | UP | H-2 class II histocompatibility antigen, A-K beta chain |
| HB2S_MOUSE | P06345 | UP | H-2 class II histocompatibility antigen, A-S beta chain |
| HB2Q_MOUSE | P06342 | UP | H-2 class II histocompatibility antigen, A-Q beta chain |
| HB2D_MOUSE | P01921 | UP | H-2 class II histocompatibility antigen, A-D beta chain |
| ASSY_MOUSE | P16460 | UP | Argininosuccinate synthase |
| TAP2_MOUSE | P36371 | UP | Antigen peptide transporter 2 |
| CO3_MOUSE | P01027 | UP | Complement C3 |
| CO4B_MOUSE | P01029 | UP | Complement C4-B |
| C4BPA_MOUSE | P08607 | UP | C4b-binding protein |
| GDIR2_MOUSE | Q61599 | UP | Rho GDP-dissociation inhibitor 2 |
| PGRP1_MOUSE | O88593 | UP | Peptidoglycan recognition protein 1 |
| NCF2_MOUSE | O70145 | UP | Neutrophil cytosol factor 2 |
| CATS_MOUSE | O70370 | UP | Cathepsin S |
| CFAB_MOUSE | P04186 | UP | Complement factor B |
| F13A_MOUSE | Q8BH61 | UP | Coagulation factor XIII A chain |
| HEMO_MOUSE | Q91X72 | UP | Hemopexin |
| ASC_MOUSE | Q9EPB4 | UP | Apoptosis-associated speck-like protein containing a CARD |
| ADPGK_MOUSE | Q8VDL4 | UP | ADP-dependent glucokinase |
| CYRIB_MOUSE | Q921M7 | UP | CYFIP-related Rac1 interactor B |
| COTL1_MOUSE | Q9CQI6 | UP | Coactosin-like protein |
| FRIH_MOUSE | P09528 | UP | Ferritin heavy chain |
| ABCB9_MOUSE | Q9JJ59 | UP | ABC-type oligopeptide transporter ABCB9 |
| COR1A_MOUSE | O89053 | UP | Coronin-1A |
| PTN6_MOUSE | P29351 | UP | Tyrosine-protein phosphatase non-receptor type 6 |
| CD44_MOUSE | P15379 | UP | CD44 antigen |
| IFM3_MOUSE | Q9CQW9 | UP | Interferon-induced transmembrane protein 3 |
| CFAH_MOUSE | P06909 | UP | Complement factor H |
| NAMPT_MOUSE | Q99KQ4 | UP | Nicotinamide phosphoribosyltransferase |
| PLSL_MOUSE | Q61233 | UP | Plastin-2 |
| CATB_MOUSE | P10605 | UP | Cathepsin B |
| FKBP5_MOUSE | Q64378 | UP | Peptidyl-prolyl cis-trans isomerase FKBP5 |
| PSB10_MOUSE | O35955 | UP | Proteasome subunit beta type-10 |
| HA1B_MOUSE | P01901 | UP | H-2 class I histocompatibility antigen, K-B alpha chain |
| HA1K_MOUSE | P04223 | UP | H-2 class I histocompatibility antigen, K-K alpha chain |
| HA11_MOUSE | P01899 | UP | H-2 class I histocompatibility antigen, D-B alpha chain |
| B2MG_MOUSE | P01887 | UP | Beta-2-microglobulin |
| NPC2_MOUSE | Q9Z0J0 | UP | NPC intracellular cholesterol transporter 2 |
| PYRG1_MOUSE | P70698 | UP | CTP synthase 1 |
| CMPK2_MOUSE | Q3U5Q7 | UP | UMP-CMP kinase 2, mitochondrial |
| GLRX1_MOUSE | Q9QUH0 | UP | Glutaredoxin-1 |
| VTNC_MOUSE | P29788 | UP | Vitronectin |
| RBM3_MOUSE | O89086 | UP | RNA-binding protein 3 |
| HMGB2_MOUSE | P30681 | UP | High mobility group protein B2 |
| IC1_MOUSE | P97290 | UP | Plasma protease C1 inhibitor |
| TCTP_MOUSE | P63028 | UP | Translationally-controlled tumor protein |
| FIBA_MOUSE | E9PV24 | UP | Fibrinogen alpha chain |
| FIBB_MOUSE | Q8K0E8 | UP | Fibrinogen beta chain |
| CERU_MOUSE | Q61147 | UP | Ceruloplasmin |
| CATZ_MOUSE | Q9WUU7 | UP | Cathepsin Z |
| **CLUS_MOUSE** | **Q06890** | **UP** | **Clusterin** |
| IPYR_MOUSE | Q9D819 | UP | Inorganic pyrophosphatase |
| LRC59_MOUSE | Q922Q8 | UP | Leucine-rich repeat-containing protein 59 |
| SAMH1_MOUSE | Q60710 | UP | Deoxynucleoside triphosphate triphosphohydrolase SAMHD1 |
| TGM2_MOUSE | P21981 | UP | Protein-glutamine gamma-glutamyltransferase 2 |
| ANXA1_MOUSE | P10107 | UP | Annexin A1 |
| AOFA_MOUSE | Q64133 | DOWN | Amine oxidase |
| AL1A1_MOUSE | P24549 | DOWN | Aldehyde dehydrogenase 1A1 |
| DHSO_MOUSE | Q64442 | DOWN | Sorbitol dehydrogenase |
| LIMC1_MOUSE | Q3UH68 | DOWN | LIM and calponin homology domains-containing protein 1 |
| TRBM_MOUSE | P15306 | DOWN | Thrombomodulin |
| PGDH_MOUSE | Q8VCC1 | DOWN | 15-hydroxyprostaglandin dehydrogenase [NAD(+)] |
| CP2F2_MOUSE | P33267 | DOWN | Cytochrome P450 2F2 |
| PXDC2_MOUSE | Q9DC11 | DOWN | Plexin domain-containing protein 2 |
| ACTC_MOUSE | P68033 | DOWN | Actin, alpha cardiac muscle 1 |
| ZYX_MOUSE | Q62523 | DOWN | Zyxin |
| LIN7C_MOUSE | O88952 | DOWN | Protein lin-7 homolog C |
| SEPT4_MOUSE | P28661 | DOWN | Septin-4 |
| ACTA_MOUSE | P62737 | DOWN | Actin, aortic smooth muscle |
| ACTH_MOUSE | P63268 | DOWN | Actin, gamma-enteric smooth muscle |
| DDAH1_MOUSE | Q9CWS0 | DOWN | N(G),N(G)-dimethylarginine dimethylaminohydrolase 1 |
| ACS2L_MOUSE | Q99NB1 | DOWN | Acetyl-coenzyme A synthetase 2-like, mitochondrial |
| AOXC_MOUSE | G3X982 | DOWN | Aldehyde oxidase 3 |
| ATPK_MOUSE | P56135 | DOWN | ATP synthase subunit f, mitochondrial |
| PYGB_MOUSE | Q8CI94 | DOWN | Glycogen phosphorylase, brain form |
| DYL1_MOUSE | P63168 | DOWN | Dynein light chain 1, cytoplasmic |
| SYNE2_MOUSE | Q6ZWQ0 | DOWN | Nesprin-2 |
| UTER_MOUSE | Q06318 | DOWN | Uteroglobin |
| CDIPT_MOUSE | Q8VDP6 | DOWN | CDP-diacylglycerol--inositol 3-phosphatidyltransferase |
| DDI2_MOUSE | A2ADY9 | DOWN | Protein DDI1 homolog 2 |
| TSN7_MOUSE | Q62283 | DOWN | Tetraspanin-7 |
| FMO3_MOUSE | P97501 | DOWN | Flavin-containing monooxygenase 3 |
| PGBM_MOUSE | Q05793 | DOWN | Basement membrane-specific heparan sulfate proteoglycan core protein |

**Supplementary Table 2.** A list of pathway analysis from the intersection data with plasma-EV and lung tissue samples.

The pathway name, raw *p*-value, and -log10 *p*-value in the AP group with the intersection data are shown. (only showed the pathway where the raw *p*-value <0.05)

| **Groups** | **Pathway name** | **Total** | **Hits** | **Raw p** | **-log10 p** | **Impact** |
| --- | --- | --- | --- | --- | --- | --- |
| AP/Sham | Arachidonic acid metabolism | 43 | 9 | 1.19E-06 | 5.9247 | 0.20552 |
| AP/Sham | Galactose metabolism | 27 | 6 | 6.25E-05 | 4.2042 | 0.42037 |
| AP/Sham | D-Amino acid metabolism | 15 | 4 | 0.00057 | 3.2438 | 0 |
| AP/Sham | Valine, leucine and isoleucine biosynthesis | 8 | 3 | 0.001023 | 2.99 | 0 |
| AP/Sham | Arginine and proline metabolism | 36 | 5 | 0.002624 | 2.581 | 0.21512 |
| AP/Sham | Valine, leucine and isoleucine degradation | 40 | 5 | 0.004216 | 2.3751 | 0.04094 |
| AP/Sham | Linoleic acid metabolism | 5 | 2 | 0.007185 | 2.1436 | 0 |
| AP/Sham | Fructose and mannose metabolism | 18 | 3 | 0.012268 | 1.9112 | 0.09765 |
| LPS/Sham | Arachidonic acid metabolism | 43 | 9 | 9.31E-09 | 8.031 | 0.20552 |
| LPS/Sham | Galactose metabolism | 27 | 6 | 2.89E-06 | 5.5394 | 0.42037 |
| LPS/Sham | Fructose and mannose metabolism | 18 | 3 | 0.002928 | 2.5335 | 0.09765 |
| LPS/Sham | Valine, leucine and isoleucine biosynthesis | 8 | 2 | 0.007177 | 2.1441 | 0 |
| LPS/Sham | Starch and sucrose metabolism | 15 | 2 | 0.025032 | 1.6015 | 0.33199 |
| LPS/Sham | Amino sugar and nucleotide sugar metabolism | 42 | 3 | 0.031501 | 1.5017 | 0 |
| LPS/Sham | Neomycin, kanamycin and gentamicin biosynthesis | 2 | 1 | 0.033406 | 1.4762 | 0 |
| COVID-19/Sham | Arachidonic acid metabolism | 43 | 12 | 4.52E-15 | 14.344 | 0.15998 |
| COVID-19/Sham | Sphingolipid metabolism | 32 | 3 | 0.007259 | 2.1391 | 0.10006 |
| COVID-19/Sham | Pyruvate metabolism | 23 | 2 | 0.034277 | 1.465 | 0.07682 |

**Supplementary Table 3.** List of all significant functions, focusing on our regulation omics data, joint p-values for all ARDS groups of lung tissue samples from OmicsNet.

Functional enrichment according to the KEGG database. The joint p-value combining metabolites and proteins is shown.

| **Pathway** | **Groups** | **HitsG** | **HitsM** | **Total hits** | **Joint p-value** |
| --- | --- | --- | --- | --- | --- |
| Glycolysis / Gluconeogenesis | AP/Sham | 8 | 6 | 14 | 1.10E-07 |
| Arachidonic acid metabolism | AP/Sham | 3 | 12 | 15 | 1.19E-07 |
| Galactose metabolism | AP/Sham | 1 | 9 | 10 | 3.52E-06 |
| Amino sugar and nucleotide sugar metabolism | AP/Sham | 4 | 11 | 15 | 4.00E-06 |
| Glucagon signaling pathway | AP/Sham | 1 | 7 | 8 | 1.13E-05 |
| Neuroactive ligand-receptor interaction | AP/Sham | 7 | 6 | 13 | 3.76E-05 |
| Platelet activation | AP/Sham | 3 | 4 | 7 | 6.45E-05 |
| Fructose and mannose metabolism | AP/Sham | 2 | 7 | 9 | 0.00013 |
| ABC transporters | AP/Sham | 7 | 9 | 16 | 0.000145 |
| Valine, leucine and isoleucine degradation | AP/Sham | 4 | 5 | 9 | 0.000311 |
| Oxytocin signaling pathway | AP/Sham | 1 | 4 | 5 | 0.000564 |
| African trypanosomiasis | AP/Sham | 1 | 3 | 4 | 0.00165 |
| Starch and sucrose metabolism | AP/Sham | 3 | 4 | 7 | 0.00177 |
| Gastric cancer | AP/Sham | 2 | 2 | 4 | 0.00191 |
| Amoebiasis | AP/Sham | 2 | 3 | 5 | 0.00208 |
| cAMP signaling pathway | AP/Sham | 2 | 4 | 6 | 0.00243 |
| Non-small cell lung cancer | AP/Sham | 2 | 2 | 4 | 0.00487 |
| Mineral absorption | AP/Sham | 1 | 4 | 5 | 0.00735 |
| Citrate cycle (TCA cycle) | AP/Sham | 4 | 2 | 6 | 0.0078 |
| Regulation of lipolysis in adipocytes | AP/Sham | 1 | 3 | 4 | 0.00791 |
| Pentose phosphate pathway | AP/Sham | 3 | 3 | 6 | 0.00975 |
| Inositol phosphate metabolism | AP/Sham | 1 | 5 | 6 | 0.0118 |
| Insulin resistance | AP/Sham | 2 | 3 | 5 | 0.0118 |
| Biosynthesis of unsaturated fatty acids | AP/Sham | 1 | 6 | 7 | 0.012 |
| Propanoate metabolism | AP/Sham | 2 | 4 | 6 | 0.0123 |
| Leishmaniasis | AP/Sham | 1 | 2 | 3 | 0.0131 |
| AMPK signaling pathway | AP/Sham | 1 | 4 | 5 | 0.0133 |
| Prolactin signaling pathway | AP/Sham | 2 | 2 | 4 | 0.0207 |
| Vascular smooth muscle contraction | AP/Sham | 2 | 2 | 4 | 0.0249 |
| Rheumatoid arthritis | AP/Sham | 3 | 1 | 4 | 0.0256 |
| Purine metabolism | AP/Sham | 7 | 4 | 11 | 0.034 |
| Fc epsilon RI signaling pathway | AP/Sham | 1 | 2 | 3 | 0.0457 |
| Antifolate resistance | AP/Sham | 1 | 2 | 3 | 0.0461 |
| Taste transduction | AP/Sham | 2 | 3 | 5 | 0.0462 |
| Pyrimidine metabolism | AP/Sham | 2 | 4 | 6 | 0.0498 |
| Platelet activation | LPS/Sham | 1 | 4 | 5 | 8.18E-05 |
| Regulation of lipolysis in adipocytes | LPS/Sham | 1 | 4 | 5 | 8.38E-05 |
| ABC transporters | LPS/Sham | 2 | 9 | 11 | 0.00018 |
| Vascular smooth muscle contraction | LPS/Sham | 5 | 2 | 7 | 0.000334 |
| Aldosterone synthesis and secretion | LPS/Sham | 1 | 4 | 5 | 0.000497 |
| Amoebiasis | LPS/Sham | 1 | 3 | 4 | 0.00113 |
| Cortisol synthesis and secretion | LPS/Sham | 1 | 3 | 4 | 0.00122 |
| Inositol phosphate metabolism | LPS/Sham | 1 | 5 | 6 | 0.00129 |
| Gastric cancer | LPS/Sham | 1 | 2 | 3 | 0.00145 |
| Starch and sucrose metabolism | LPS/Sham | 1 | 4 | 5 | 0.00217 |
| Neuroactive ligand-receptor interaction | LPS/Sham | 1 | 5 | 6 | 0.00273 |
| Aminoacyl-tRNA biosynthesis | LPS/Sham | 1 | 4 | 5 | 0.0108 |
| Pentose phosphate pathway | LPS/Sham | 1 | 3 | 4 | 0.0129 |
| Purine metabolism | LPS/Sham | 4 | 4 | 8 | 0.0132 |
| Caffeine metabolism | LPS/Sham | 1 | 2 | 3 | 0.0171 |
| Biosynthesis of unsaturated fatty acids | LPS/Sham | 1 | 4 | 5 | 0.0247 |
| Endocrine resistance | LPS/Sham | 2 | 1 | 3 | 0.037 |
| Rheumatoid arthritis | LPS/Sham | 1 | 1 | 2 | 0.0421 |
| Salmonella infection | LPS/Sham | 1 | 1 | 2 | 0.0496 |
| Gastric cancer | COVID-19/Sham | 1 | 2 | 3 | 6.74E-05 |
| Salmonella infection | COVID-19/Sham | 1 | 1 | 2 | 0.00962 |
| Vascular smooth muscle contraction | COVID-19/Sham | 2 | 1 | 3 | 0.0111 |
| Th17 cell differentiation | COVID-19/Sham | 1 | 1 | 2 | 0.0153 |
| Amoebiasis | COVID-19/Sham | 1 | 1 | 2 | 0.0329 |
| Platelet activation | COVID-19/Sham | 1 | 1 | 2 | 0.0362 |
| Cortisol synthesis and secretion | COVID-19/Sham | 1 | 1 | 2 | 0.0363 |
| ABC transporters | COVID-19/Sham | 2 | 2 | 4 | 0.0384 |
| Arginine biosynthesis | COVID-19/Sham | 1 | 1 | 2 | 0.0389 |
| Gap junction | COVID-19/Sham | 1 | 1 | 2 | 0.0451 |

HitsG: Gene name from protein hit number.

HitsM: Metabolite hit number.

**Supplementary Table 4.** A list of DEMs' intersection data with total six types of plasma-EV (pEV) and lung tissue samples.

| **Metabolites** | **Description** | **Formula** | **Electrode**  **(pEV)** | **m/z from pEV** | **Electrode**  **(lung tissue)** | **m/z from lung tissue**  **(LPS and COVID group)** | **m/z from lung tissue**  **(only AP group)** |
| --- | --- | --- | --- | --- | --- | --- | --- |
| HMDB0260512 | MG(20:3(6,8,11)-OH(5)/0:0/0:0) | C_23_H_40_O_5_ | M+Na | 14.95_419.2760m/z | M+H | 9.34_397.2957m/z | 9.44_419.2777m/z |
| HMDB0260564 | MG(0:0/20:3(6,8,11)-OH(5)/0:0) | C_23_H_40_O_5_ | M+Na | 14.95_419.2760m/z | M+H | 9.34_397.2957m/z | 9.44_419.2777m/z |

| **Metabolites** | **Description** | **Formula** | **Electrode**  **(pEV)** | **m/z from pEV** | **Electrode**  **(lung tissue)** | **m/z from lung tissue**  **(AP and COVID group)** | **m/z from lung tissue**  **(AP and LPS group)** |
| --- | --- | --- | --- | --- | --- | --- | --- |
| HMDB0031415 | 2,6-Dimethyl-2,4-heptadiene | C_9_H_16_ | M+H | 12.13_125.1331m/z | M+H | 9.01_125.1330m/z | 12.12_125.1331m/z |
| HMDB0037777 | 2-Isopropyl-1,4-hexadiene | C_9_H_16_ | M+H | 12.13_125.1331m/z | M+H | 9.01_125.1330m/z | 12.12_125.1331m/z |
| HMDB0061920 | Prop-2-enylcyclohexane | C_9_H_16_ | M+H | 12.13_125.1331m/z | M+H | 9.01_125.1330m/z | 12.12_125.1331m/z |
| HMDB0244025 | 1,1-Dimethyl-2-(2-methyl-1-propenyl) cyclopropane | C_9_H_16_ | M+H | 12.13_125.1331m/z | M+H | 9.01_125.1330m/z | 12.12_125.1331m/z |
| HMDB0253139 | Hexahydroindan | C_9_H_16_ | M+H | 12.13_125.1331m/z | M+H | 9.01_125.1330m/z | 12.12_125.1331m/z |

| **Metabolites** | **Description** | **Formula** | **Electrode**  **(pEV)** | **m/z from pEV** | **Electrode**  **(lung tissue)** | **m/z from lung tissue** |
| --- | --- | --- | --- | --- | --- | --- |
| HMDB0000451 | cis-4-Hydroxycyclohexylacetic acid | C_8_H_14_O_3_ | M+2H | 6.43_80.0547m/z | M+2H | 9.17_80.0546m/z |
| HMDB0000909 | trans-4-Hydroxycyclohexylacetic acid | C_8_H_14_O_3_ | M+2H | 6.43_80.0547m/z | M+2H | 9.17_80.0546m/z |
| HMDB0010721 | 3-Oxooctanoic acid | C_8_H_14_O_3_ | M+2H | 6.43_80.0547m/z | M+2H | 9.17_80.0546m/z |
| HMDB0013211 | Alpha-Ketooctanoic acid | C_8_H_14_O_3_ | M+2H | 6.43_80.0547m/z | M+2H | 9.17_80.0546m/z |
| HMDB0030303 | 6-Ethyl-1-methyl-2,7,8-trioxabicyclo [3.2.1] octane | C_8_H_14_O_3_ | M+2H | 6.43_80.0547m/z | M+2H | 9.17_80.0546m/z |
| HMDB0031177 | Tetrahydrofurfuryl propionate | C_8_H_14_O_3_ | M+2H | 6.43_80.0547m/z | M+2H | 9.17_80.0546m/z |
| HMDB0031307 | Ethyl 3-oxohexanoate | C_8_H_14_O_3_ | M+2H | 6.43_80.0547m/z | M+2H | 9.17_80.0546m/z |
| HMDB0036230 | 1-Methyl-2-oxopropyl butyrate | C_8_H_14_O_3_ | M+2H | 6.43_80.0547m/z | M+2H | 9.17_80.0546m/z |
| HMDB0036395 | 2-Methylpropyl 3-oxobutanoate | C_8_H_14_O_3_ | M+2H | 6.43_80.0547m/z | M+2H | 9.17_80.0546m/z |
| HMDB0038305 | 3-Methylbutyl 2-oxopropanoate | C_8_H_14_O_3_ | M+2H | 6.43_80.0547m/z | M+2H | 9.17_80.0546m/z |
| HMDB0040447 | Butyl acetoacetate | C_8_H_14_O_3_ | M+2H | 6.43_80.0547m/z | M+2H | 9.17_80.0546m/z |
| HMDB0041616 | Propyl levulinate | C_8_H_14_O_3_ | M+2H | 6.43_80.0547m/z | M+2H | 9.17_80.0546m/z |
| HMDB0059938 | 5-Butyl-1,4-dioxan-2-one | C_8_H_14_O_3_ | M+2H | 6.43_80.0547m/z | M+2H | 9.17_80.0546m/z |
| HMDB0059939 | 6-Butyl-1,4-dioxan-2-one | C_8_H_14_O_3_ | M+2H | 6.43_80.0547m/z | M+2H | 9.17_80.0546m/z |
| HMDB0060683 | 2-n-Propyl-4-oxopentanoic acid | C_8_H_14_O_3_ | M+2H | 6.43_80.0547m/z | M+2H | 9.17_80.0546m/z |
| HMDB0060685 | 3-Oxovalproic acid | C_8_H_14_O_3_ | M+2H | 6.43_80.0547m/z | M+2H | 9.17_80.0546m/z |
| HMDB0062788 | Isobutyric Acid Anhydride | C_8_H_14_O_3_ | M+2H | 6.43_80.0547m/z | M+2H | 9.17_80.0546m/z |
| HMDB0303914 | Butanoic anhydride | C_8_H_14_O_3_ | M+2H | 6.43_80.0547m/z | M+2H | 9.17_80.0546m/z |
| HMDB0013336 | 3-Hydroxyhexadecanoylcarnitine | C_23_H_45_NO_5_ | M-H | 9.43_414.3218m/z | M+K | 8.70_454.2949m/z |
| HMDB0061642 | 3-hydroxyhexadecanoyl carnitine | C_23_H_45_NO_5_ | M-H | 9.43_414.3218m/z | M+K | 8.70_454.2949m/z |
| HMDB0241458 | 16-Hydroxyhexadecanoylcarnitine | C_23_H_45_NO_5_ | M-H | 9.43_414.3218m/z | M+K | 8.70_454.2949m/z |
| HMDB0241459 | (2S)-2-Hydroxyhexadecanoylcarnitine | C_23_H_45_NO_5_ | M-H | 9.43_414.3218m/z | M+K | 8.70_454.2949m/z |
| HMDB0241460 | 5-Hydroxyhexadecanoylcarnitine | C_23_H_45_NO_5_ | M-H | 9.43_414.3218m/z | M+K | 8.70_454.2949m/z |
| HMDB0241461 | 7-Hydroxyhexadecanoylcarnitine | C_23_H_45_NO_5_ | M-H | 9.43_414.3218m/z | M+K | 8.70_454.2949m/z |
| HMDB0241462 | 8-Hydroxyhexadecanoylcarnitine | C_23_H_45_NO_5_ | M-H | 9.43_414.3218m/z | M+K | 8.70_454.2949m/z |
| HMDB0241463 | 9-Hydroxyhexadecanoylcarnitine | C_23_H_45_NO_5_ | M-H | 9.43_414.3218m/z | M+K | 8.70_454.2949m/z |
| HMDB0241464 | 10-Hydroxyhexadecanoylcarnitine | C_23_H_45_NO_5_ | M-H | 9.43_414.3218m/z | M+K | 8.70_454.2949m/z |
| HMDB0241465 | 11-Hydroxyhexadecanoylcarnitine | C_23_H_45_NO_5_ | M-H | 9.43_414.3218m/z | M+K | 8.70_454.2949m/z |
| HMDB0241466 | 12-Hydroxyhexadecanoylcarnitine | C_23_H_45_NO_5_ | M-H | 9.43_414.3218m/z | M+K | 8.70_454.2949m/z |
| HMDB0241467 | 13-Hydroxyhexadecanoylcarnitine | C_23_H_45_NO_5_ | M-H | 9.43_414.3218m/z | M+K | 8.70_454.2949m/z |
| HMDB0241468 | 6-Hydroxyhexadecanoylcarnitine | C_23_H_45_NO_5_ | M-H | 9.43_414.3218m/z | M+K | 8.70_454.2949m/z |
| HMDB0001220 | Prostaglandin E2 | C_20_H_32_O_5_ | M-H | 8.82_351.2171m/z | M-H | 7.89_351.2160m/z |
| HMDB0001320 | (13E)-11a-Hydroxy-9,15-dioxoprost-13-enoic acid | C_20_H_32_O_5_ | M-H | 8.82_351.2171m/z | M-H | 7.89_351.2160m/z |
| HMDB0001335 | Prostaglandin I2 | C_20_H_32_O_5_ | M-H | 8.82_351.2171m/z | M-H | 7.89_351.2160m/z |
| HMDB0001381 | Prostaglandin H2 | C_20_H_32_O_5_ | M-H | 8.82_351.2171m/z | M-H | 7.89_351.2160m/z |
| HMDB0001403 | Prostaglandin D2 | C_20_H_32_O_5_ | M-H | 8.82_351.2171m/z | M-H | 7.89_351.2160m/z |
| HMDB0001452 | Thromboxane A2 | C_20_H_32_O_5_ | M-H | 8.82_351.2171m/z | M-H | 7.89_351.2160m/z |
| HMDB0001509 | 20-Hydroxy-leukotriene B4 | C_20_H_32_O_5_ | M-H | 8.82_351.2171m/z | M-H | 7.89_351.2160m/z |
| HMDB0002122 | Prostaglandin F3a | C_20_H_32_O_5_ | M-H | 8.82_351.2171m/z | M-H | 7.89_351.2160m/z |
| HMDB0002132 | 8-iso-PGF3a | C_20_H_32_O_5_ | M-H | 8.82_351.2171m/z | M-H | 7.89_351.2160m/z |
| HMDB0002363 | Levuglandin E2 | C_20_H_32_O_5_ | M-H | 8.82_351.2171m/z | M-H | 7.89_351.2160m/z |
| HMDB0002400 | Levuglandin D2 | C_20_H_32_O_5_ | M-H | 8.82_351.2171m/z | M-H | 7.89_351.2160m/z |
| HMDB0002776 | 13,14-Dihydro-15-keto-PGE2 | C_20_H_32_O_5_ | M-H | 8.82_351.2171m/z | M-H | 7.89_351.2160m/z |
| HMDB0004240 | 15-Keto-prostaglandin F2a | C_20_H_32_O_5_ | M-H | 8.82_351.2171m/z | M-H | 7.89_351.2160m/z |
| HMDB0004385 | Lipoxin A4 | C_20_H_32_O_5_ | M-H | 8.82_351.2171m/z | M-H | 7.89_351.2160m/z |
| HMDB0005077 | 8-iso-15-keto-PGF2a | C_20_H_32_O_5_ | M-H | 8.82_351.2171m/z | M-H | 7.89_351.2160m/z |
| HMDB0005082 | Lipoxin B4 | C_20_H_32_O_5_ | M-H | 8.82_351.2171m/z | M-H | 7.89_351.2160m/z |
| HMDB0005844 | 8-isoprostaglandin E2 | C_20_H_32_O_5_ | M-H | 8.82_351.2171m/z | M-H | 7.89_351.2160m/z |
| HMDB0012481 | (5Z)-(15S)-11alpha-Hydroxy-9,15-dioxoprostanoate | C_20_H_32_O_5_ | M-H | 8.82_351.2171m/z | M-H | 7.89_351.2160m/z |
| HMDB0012564 | 13,14-Dihydro-15-oxo-lipoxin A4 | C_20_H_32_O_5_ | M-H | 8.82_351.2171m/z | M-H | 7.89_351.2160m/z |
| HMDB0012587 | 15-Epi-lipoxin A4 | C_20_H_32_O_5_ | M-H | 8.82_351.2171m/z | M-H | 7.89_351.2160m/z |
| HMDB0034676 | Cinncassiol D1 | C_20_H_32_O_5_ | M-H | 8.82_351.2171m/z | M-H | 7.89_351.2160m/z |
| HMDB0035064 | Cinncassiol D4 | C_20_H_32_O_5_ | M-H | 8.82_351.2171m/z | M-H | 7.89_351.2160m/z |
| HMDB0035338 | Sterebin B | C_20_H_32_O_5_ | M-H | 8.82_351.2171m/z | M-H | 7.89_351.2160m/z |
| HMDB0035339 | Sterebin C | C_20_H_32_O_5_ | M-H | 8.82_351.2171m/z | M-H | 7.89_351.2160m/z |
| HMDB0036756 | (ent-6alpha,7alpha,16alphaH)-6,7,17-Trihydroxy-19-kauranoic acid | C_20_H_32_O_5_ | M-H | 8.82_351.2171m/z | M-H | 7.89_351.2160m/z |
| HMDB0060041 | 11b-PGE2 | C_20_H_32_O_5_ | M-H | 8.82_351.2171m/z | M-H | 7.89_351.2160m/z |
| HMDB0060042 | 13,14-Dihydro-15-keto-PGD2 | C_20_H_32_O_5_ | M-H | 8.82_351.2171m/z | M-H | 7.89_351.2160m/z |
| HMDB0062291 | 15-oxo-5S,6R-dihydroxy-7E,9E,11Z-eicosatrienoic acid | C_20_H_32_O_5_ | M-H | 8.82_351.2171m/z | M-H | 7.89_351.2160m/z |
| HMDB0062691 | 15-dehydro-prostaglandin E1(1-) | C_20_H_32_O_5_ | M-H | 8.82_351.2171m/z | M-H | 7.89_351.2160m/z |
| HMDB0062798 | 5-hydroperoxy-15-HETE | C_20_H_32_O_5_ | M-H | 8.82_351.2171m/z | M-H | 7.89_351.2160m/z |
| HMDB0244513 | (Z)-7-((1R,5S)-5-Hydroxy-3-oxo-2-(3-oxooctyl)cyclopentyl)hept-5-enoic acid | C_20_H_32_O_5_ | M-H | 8.82_351.2171m/z | M-H | 7.89_351.2160m/z |
| HMDB0245567 | 20-hydroxyleukotriene B4 | C_20_H_32_O_5_ | M-H | 8.82_351.2171m/z | M-H | 7.89_351.2160m/z |
| HMDB0246193 | Prostacyclin (not favourable) | C_20_H_32_O_5_ | M-H | 8.82_351.2171m/z | M-H | 7.89_351.2160m/z |
| HMDB0247662 | (5S,6E,8E,10E,12E,14S,15R)-5,14,15-Trihydroxyicosa-6,8,10,12-tetraenoic acid | C_20_H_32_O_5_ | M-H | 8.82_351.2171m/z | M-H | 7.89_351.2160m/z |
| HMDB0247796 | 7-[(1R,2R,3R)-3-Hydroxy-2-[(3S)-3-hydroxyoct-1-enyl]-5-oxocyclopentyl]-5-heptenoic acid | C_20_H_32_O_5_ | M-H | 8.82_351.2171m/z | M-H | 7.89_351.2160m/z |
| HMDB0251839 | Epi-Lipoxin A4 | C_20_H_32_O_5_ | M-H | 8.82_351.2171m/z | M-H | 7.89_351.2160m/z |
| HMDB0302207 | Prostaglandin E-2 | C_20_H_32_O_5_ | M-H | 8.82_351.2171m/z | M-H | 7.89_351.2160m/z |
| HMDB0303098 | [6]-Gingerdiol acetate methyl ether | C_20_H_32_O_5_ | M-H | 8.82_351.2171m/z | M-H | 7.89_351.2160m/z |
| HMDB0245262 | 2-Nitrophenyl octyl ether | C_14_H_21_NO_3_ | M-H | 11.20_250.1450m/z | M+H | 13.97_252.1597m/z |
